# A simple model of a quadruped discovers single-foot walking and trotting as energy optimal strategies

**DOI:** 10.1101/580779

**Authors:** Delyle T. Polet, John E. A. Bertram

## Abstract

It is widely held that quadrupeds choose steady gaits that minimize their energetic cost of transport, but it is difficult to explore the entire range of possible footfall sequences empirically. We present a simple model of a quadruped that can spontaneously produce any of the 2300+ planar footfall sequences available to quadrupeds. Through trajectory optimization of a work and force-rate cost, and a large sample of random initial guesses, we provide stronger evidence towards the global optimality of symmetrical four-beat walking at low speeds and two beat running (trotting) at intermediate speeds. Using input parameters based on measurements in dogs (*Canis lupus familiaris*), the model predicts the correct phase offset in walking and a realistic walk-trot transition speed. It also spontaneously reproduces the double-hump ground reaction force profile observed in walking, and the smooth single-hump profile observed in trotting, despite the model’s lack of springs. However, the model incorrectly predicts duty factor at the slowest speeds, and does not choose galloping as globally optimal at high speeds. These limitations might point to missing levels of complexity that could be added to future quadrupedal models, but the present results suggest that massive legs, elastic components, head-neck oscillations and even multi-linked legs are not critical determiners for the optimality of natural quadrupedal walking and trotting.

**Author summary:** Why do quadrupedal mammals move in consistent ways, when so many options are available? We tackled this problem by determining energetically-optimal gaits using a simple computational model of a four-legged animal. The model can use virtually any pattern of movement (physics-permitting!) but selects natural movement strategies as energetically optimal. The similarities between the computer-based predictions and natural animal movement are striking, and suggest mammals utilize movement strategies that optimize energy use when they move.

## Introduction

When unconstrained at a given speed, members of a species of quadrupedal will generally select a common gait, which is seldom unique to that species alone [1–4]. The consistency of gait choice is remarkable, given how many alternatives exist. McGhee and Jain [5] noted that while 5040 unique footfall sequences are available to quadrupeds, only 21 varieties had been reported by the mid-1970s. Within these varieties, some very consistent patterns emerge. With few exceptions, mammals choose a lateral or diagonal sequence (four-beat) walk at slow speeds, a running trot (two-beat) at intermediate speeds, and a gallop at high speeds [6, 7]. Before the trot-gallop transition, healthy mammals use highly symmetrical gaits, with foot touchdown and liftoff on the left side of the body being repeated exactly half a stride later on the right side [1, 8, 9]. Why should the same patterns repeatedly emerge among quadrupedal mammals, despite the incredible morphological diversity and near limitless gait possibilities?

It is widely held that the gaits chosen by quadrupeds at a particular speed are those that minimize metabolic energy expenditure [1–4, 10]. Since locomotion is an energy-intensive activity, it is reasonable to suppose that animals have evolved some means to lower the energetic cost of locomotion. Indeed, humans seem remarkably sensitive to cost of transport, or energetic expenditure per unit weight per unit distance, even shifting their gait in response to unusual stimuli in accordance with energetic cost mitigation [11–13]. It is more difficult to train animals to perform unusual gait tasks, but Hoyt and Taylor [10] showed that horses will exhibit increased metabolic cost of transport if they perform walking, trotting or galloping outside their natural speed ranges, and tend to chose speeds that minimize cost of transport within each gait.

But why are particular gaits energy minimizing for particular animals at particular speeds? Some have pointed to elastic elements as a key aspect [14–16]. Gan *et al.* [15] found that natural gaits of horses could be replicated by a passive dynamic model with springs as legs. Xi *et al.* [16] found that a four-beat walk is work minimizing at slow speeds and a two-beat run (trotting or pacing) minimizes work at high speeds. They pointed to their use of elastic elements as an important reason why their model exhibited a double-hump ground reaction force (GRF) profile in walking and single hump GRFs in running

If elastic elements are prerequisites for gait optimality, it has important implications for the evolutionary history of gait. In particular, it suggests that appropriately-tuned tendons must have evolved *before* the gaits we see in modern organisms first appeared in their ancestors [17].

We may do well in taking guidance from bipedal research using work-based optimization methods. By incrementally increasing the complexity of the models used, we can determine what parameters are necessary prerequisites for generating the phenomenon of interest. Srinivasan and Ruina [18] used a “minimal” biped; effectively, a point mass with one massless leg. (As it was constrained to perform symmetrical gaits with no double-stance, the single leg could be reflected into the other leg for the next step, effectively representing a biped.) By minimizing work at a given step length and speed, the model predicted pendular walking at slow speeds and running at high speeds. However, it could not recreate double-stance dynamics, had infinitesimal stance duration in running, and discovered a strange “pendular run” at some combinations of speed and stride length.

Hasaneini *et al.* [19] added a trunk and legs with realistic masses and moments. Their model discovered natural walking and running, as well as forward leaning of the torso in vertical ascent, and bang-coast-bang swing dynamics among other commonly-reported features of natural human gait. Importantly, neither model used elastic elements, and both discovered double-hump GRFs in walking and single-hump in running, suggesting that elastic morphological features are not necessary prerequisites to the optimality of the solutions observed (and more likely, evolutionary post-adaptations).

In complex models of locomotion, the solution space of possible gaits tends to increase in scope and dimensionality, and so is necessarily more difficult to explore. Xi *et al.* [16] relied on a small number of preset footfall sequences to seed optimization, bringing into question whether other footfall sequences not explored in the study might have been energy-minimizing.

An advantage of the model in [16] was an explicit modelling of rate-dependent energetic penalties; in their case viscous damping. While limb work was the only cost in the above bipedal models, there is increasing evidence from the human physiology literature that muscle activation costs, in addition to muscular work, are an important determiner of gait. That is, there is increased metabolic cost for changing muscular forces quickly, even if work is held constant [20–22]. While the mechanism is currently unknown, rate penalties explain much about gait selection, particularly the stride length and duty factor vs. speed relationship [23–25].

To what extent can a simple, minimally constrained energy-optimization model, without springs, forced pendular exchange or forced stability, predict gait choice in quadrupeds? Do natural gaits emerge from such a simplified model and are they energy minimizing? Does a force-rate cost substantially enhance empirical agreement? We explore these questions with a planar, two-point-mass optimization model that can perform any of the footfall sequences available to quadrupeds. We compare the simulation results to empirical data on dogs only, but we anticipate the results are more broadly applicable across quadrupedal mammals.

## Methods

### Trajectory optimization of a minimal quadruped

#### Generalized model of a quadruped

We use a simplified model of a quadruped based on the bipedal model in [18], but extended to four legs. A rigid trunk connects two point masses. Each point mass has two massless limbs that can extend or contract. The quadruped is two-dimensional and lives in the sagittal plane (Fig 1).

**Fig 1.**
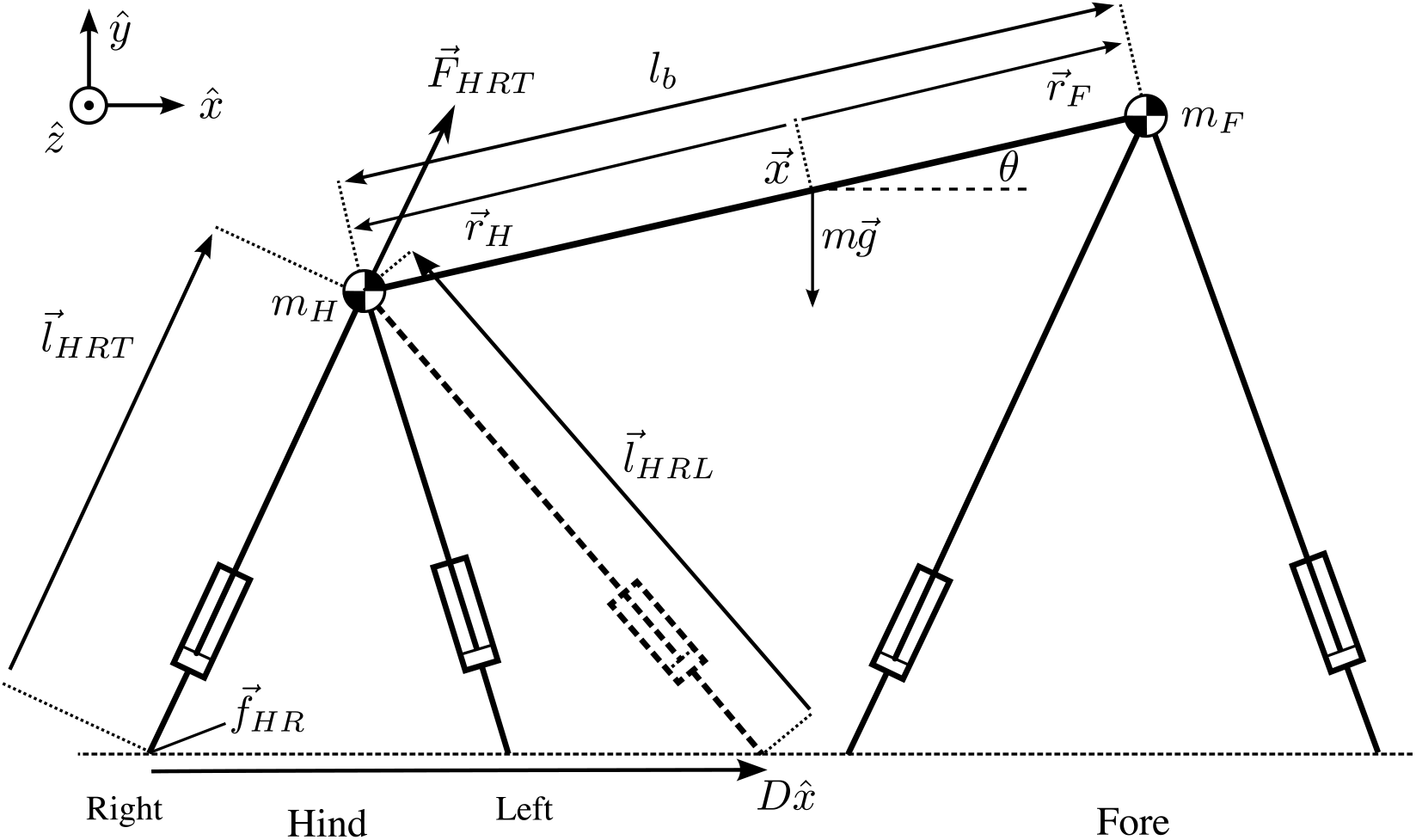
The simple quadrupedal model used in this paper. Two point masses sit on massless legs that can extend and contract. The point masses are connected with a rigid trunk. Five morphological parameters are necessary as input: the mass of the fore and hind quarters (*m*_*F*_ and *m*_*H*_, respectively), the trunk length (*l*_*b*_), and the maximum length of the fore and hind limbs (*l*_Fmax_ and *l*_Hmax_, respectively). Two kinematic variables are also necessary: the stride length (*D*) and the average horizontal speed (*U*). From these seven constant inputs, the optimal control chooses footfall positions (*f*_*ij*_) and over twenty independent temporal variables, including ground reaction forces and their rate of change (*F*_*ijk*_ and 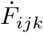), center of mass position and velocity (**x** and 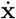) and body pitch angle and angular velocity (*θ* and 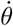).

**Fig 2.**
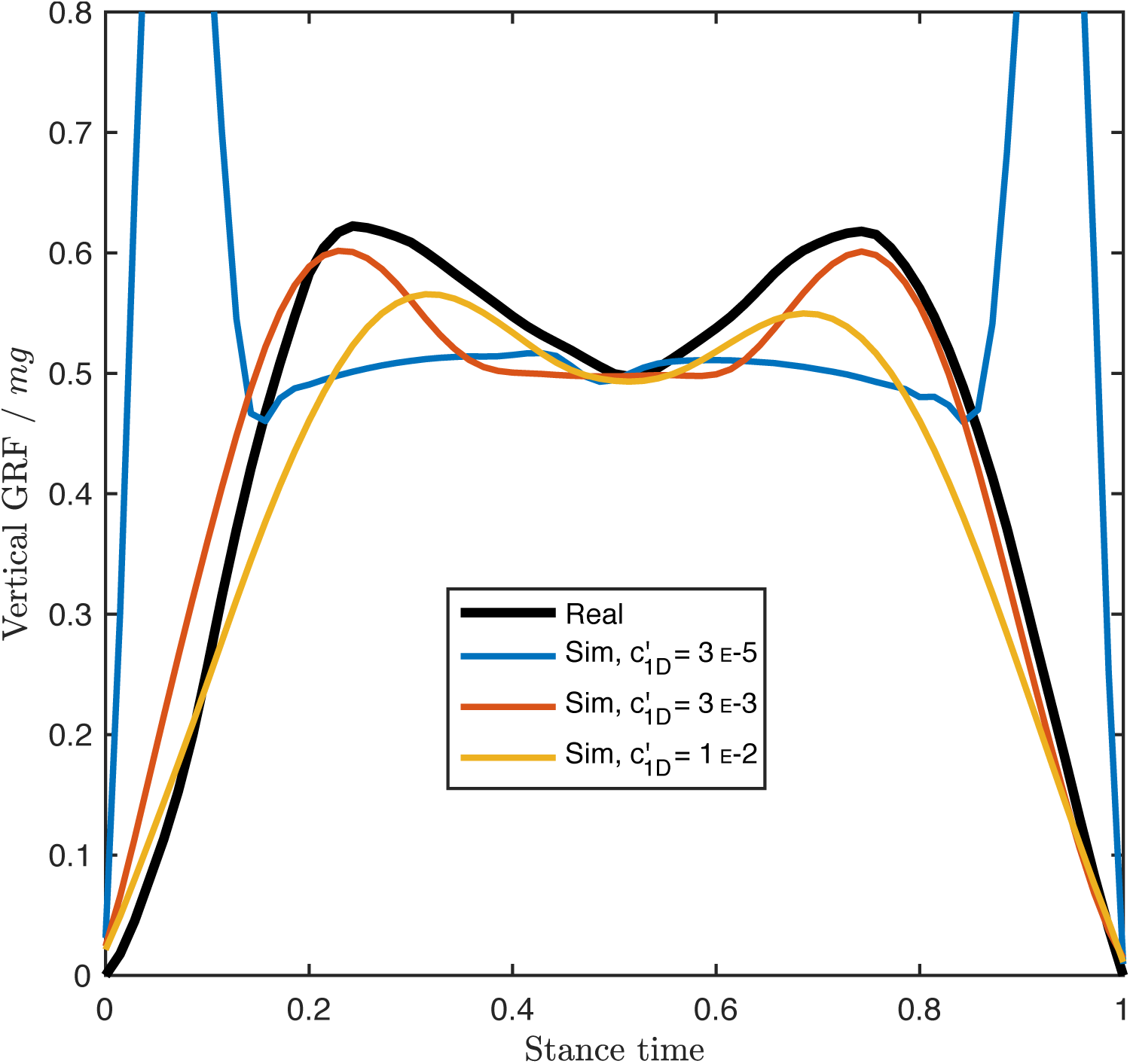
Empirical forelimb GRF (black) from [33] compared to model predictions for a range of force-rate penalty coefficients (*c*_1_ in Eq 2). At lower 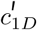, the solution becomes more impulsive, as expected from a work-minimizing “bang-coast-bang” solution [30]. As 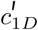 increases, the force peaks become more shallow, but in all cases the optimal solution maintains the double-hump profile characteristic of walking bipeds and quadrupeds under most situations. The morphological data is based on a Dalmation from [33] (see also Fig 4B)

Each limb comes as a “trailing” and “leading” virtual limb pair. A “trailing” limb acts through a rear footfall position, and a “leading” limb acts through a forward footfall position exactly one stride length (*D*) in advance of the trailing footfall position (the leading right-hind in Fig 1 is shown as a dotted line). The trailing limb is active *before* the leading limb (following convention, *trailing* and *leading* are spatial terms, not temporal ones). Since we are interested in periodic motion, we can always shift the temporal frame of reference so that, for one limb, touchdown occurs before liftoff. For this limb, only one virtual limb and one footfall position is necessary.

The organism travels in a vertical gravitational field with downward acceleration *g* at an average horizontal speed *U* in stride length *D*. The quadruped has forequarter mass *m*_*F*_ and hindquarter mass *m*_*H*_. Its maximum limb lengths are *l*_Fmax_ and *l*_Hmax_ for the fore and hindlimbs, respectively.

We use trajectory optimization techniques (sometimes referred-to as optimal control) to discover optimal gaits (for a beginner’s overview see [26]; for a more advanced treatment see [27]). The basic idea is that the aspects of gait the organism can directly control (*e.g.* footfall positions and forces in time) are chosen so as to minimize some cost function within particular constraints. These constraints can include physical or geometric limitations (*e.g.* mechanical dynamics or maximum limb length) or even biological limitations (*e.g.* an upper bound on force output or muscle shortening rate).

Our hypothesis is that quadrupedal mammals choose whichever gait minimizes metabolic cost per unit distance. The primary contributor to energetic cost is limb work, the combined positive and negative work performed by each limb:

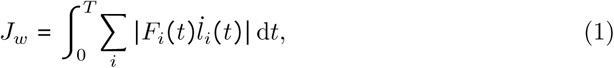

where *T* is the stride period, *F*_*i*_ is the absolute force acting through limb *i* and *l*_*i*_ is the length of limb *i. l*_*i*_ is defined as the distance between the fore-or hindquarters and footfall position *f*_*i*_ (see S1 Appendix). Note that we make no distinction between positive and negative muscle work efficiencies. While these efficiencies can alter optimal solutions in unsteady gaits [28], in steady gaits on level ground an equal amount of positive and negative work must be performed. Therefore, in the absence of other forms of dissipation (like foot-ground collisions or viscous effects) or asymmetric elastic energy recovery, the relative efficiencies of positive and negative work do not change the optimal solution.

The secondary cost is that of a force-rate penalty. Since the exact choice of rate penalty has little effect on the agreement with empirical data [29] (some fitting is required in any case, since the mechanism is unknown), we used a well-behaved squared cost in force rate,

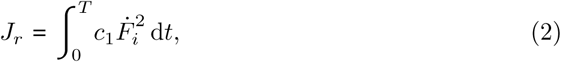

where *c*_1_ is a fitting parameter. A side benefit of the force rate cost is that it smooths the otherwise impulsive solutions that result from work-minimizing solutions [29, 30].

The resulting objective of the optimization (in the absence of augmentation for complementarity minimization), reflecting the metabolic cost of transport in the absence of a basal metabolic rate, is

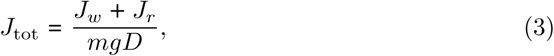

where *D* is the stride length.

#### Variables

Trajectory optimization involves two classes of time-dependent variables. Any variable that does not have a time derivative constraint is a “control variable”. For the ideal problem, these are the time derivative of limb forces, 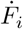. Those which have time derivative constraints are called “state variables”. The limb forces *F*_*i*_ are one such state, as are the positional variables *x* and *y* (the horizontal and vertical position of the center of mass), and *θ*, the trunk pitch angle (Fig 1). Since the dynamics are constraints on acceleration, the time derivatives of *x, y* and *θ* are also states. Finally, “parameters” are time-invariant quantities that are unknown at the outset of the problem. For the quadrupedal locomotion problem, these are the footfall positions *f*_*i*_ (Fig 1).

#### Constraints

Single-phase optimal control problems involve four types of constraints: Path and Endpoint Constraints and Variable and Integral Bounds. Path constraints must be satisfied for all time, whereas Endpoint Constraints must be satisfied at the temporal start and end of the problem. Variable bounds are upper and/or lower limits to the magnitude of variables that must also be satisfied for all time, whereas Integral bounds are limits to the value of mathematical statements integrated through time.

The mathematical statements of the constraints of the problem are stated in S1 Appendix. Briefly, solutions to the optimal control problem must satisfy the following path constraints:

P1. mechanical dynamics are satisfied,

P2. fore and hind masses must always be above ground,

P3. limb lengths are not to exceed prescribed values (measured limb length),

P4. leading limbs are active strictly after trailing limbs, and

P5. limbs are below the body: forelimb torques are always positive and hindlimb torques are always negative.

Constraint P3 determines the stepping sequence. In effect, all limbs are in contact with the ground at all times; however, the forces may only be active when limbs are sufficiently short. Once a solution is found, we can examine the ground reaction forces to assess contact. If force is nonzero, that limb is in contact with the ground; if forces are zero, that limb is not in contact. With this method, we do not need to specify contact sequence *a priori*, an advantage over the multiple shooting method used by [16]. One disadvantage of this method is that solutions exhibit a discontinuity at ground contact, which requires special treatment and makes the problem more difficult to solve. The relative simplicity of our optimal control problem makes the implementation tractable, despite this disadvantage.

Constraint P5 ensures that legs are always ventral to the body, a sensible limitation for cursorial quadrupedal mammals. Curiously, if this constraint was removed, the simulation frequently discovered bipedal running. This finding deserves further analysis as it may have implications for the evolution of bipedal locomotion, but is beyond the scope of the present study.

The following endpoint constraints are prescribed:

E1. kinematics are periodic, and

E2. Force for each given limb is equal at the start and end of stride (Force Continuity) Both endpoint constraints come out of the definition of a steady gait. An integral constraint is also prescribed:

I1. the vertical and fore-aft impulse generated by the left legs is equal to the impulse generated by the right legs.

The rationale is that breaking this constraint would cause the animal to turn or tip. Including the constraint lowered solve time, while removing the constraint did not seem to change the optimal solution.

Variables in optimal control problems may be bounded in some way. For the present case, the ideal bounds on variables and time are:

B1. Ground reaction forces are pushing forces 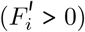.

B2. Reference limb is not active at boundary 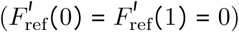.

B3. The horizontal position of the center of mass starts at *x* = 0 and ends at *x* = *D*.

B4. The simulation starts at *t* = 0 and ends at *t* = *U / D*,

where *U* is the mean fore-aft speed during a stride. Altogether, this idealized problem involves 27 unknown variables: 16 state variables, 7 control variables, and four parameters. These unknowns are all determined by optimizing the objective while satisfying the constraints. Only five morphological parameters are needed as inputs: the body length, fore-and hindlimb lengths, and mass of the fore-and hindquarters. Three additional inputs are required: Average horizontal speed, stride length, and gravitational acceleration.

However, there are several numerical issues with the above implementation. These are resolved by alterations to the optimal control problem, as discussed in the next section

#### Numerically well-conditioned optimal control problem

Three numerical issues arise in the ideal implementation. The first is the presence of a non-smooth objective due to the absolute-value function of the integrand (Eq 1). The second is due to non-smooth optimal solutions. The final issue is due to poor scaling, arising because several variables have vastly different magnitudes. Each issue can be ameliorated by reformulating the problem.

Non-smooth functions in optimal control problems can be reformulated using slack variables [27]. Two slack variables are required for every instance of the absolute value function; thus 14 new controls are introduced (*p*_*i*_ ≥ 0 and *q*_*i*_ ≥ 0). In addition, new path constraints are required, one set for redefining the operand of the absolute value function (S1 Appendix), and another for enforcing orthogonality of the slack variables:

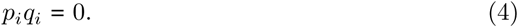

Unfortunately, Eq 4 is a complementarity condition, as are several other constraints (P3, P4 and P5) which require special treatment [27, 31]. Complementarity constraints are of the form,

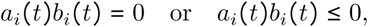

where *a*_*i*_(*t*) and *b*_*i*_(*t*) are arbitrary variables [27]. One way to manage these conditions is to introduce “relaxation parameters” that effectively smooth the constraint, as

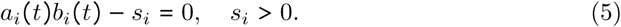

These relaxation parameters decrease the accuracy of the solution (compared to the solution of the non-smoothed problem), but by iteratively reducing the values of *s*_*i*_, we can get arbitrarily close to the true solution.

We can further ratchet down the relaxation parameters by incorporating them into the optimization problem. We make *s*_*i*_ decision variables (controls), and augment them with the penalty

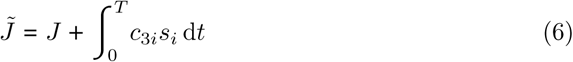

where *c*_3*i*_ is a constant chosen to be sufficiently large [31]. Then, the optimizer tends to reduce the value of *s*_*i*_, making the solution more accurate.

While it is not necessary to use both the strategy 5 and 6, we found that incorporating both reduced solve time and eased implementation. Our strategy was to start by not enforcing complementarity conditions (*c*_3*i*_ were 0 and *s*_*i*_ were allowed to take on any positive value) and then to increased the coefficients *c*_3*i*_ as we increased the resolution on each iteration. An exception to this was in the slack variable complementarity condition itself (Eq 4), which, on every iteration except the first, was not enforced except through the augmentation

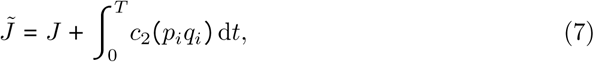

where *c*_2_ = 1 E - 3.

The resulting cost function for the optimization (prior to normalization), was

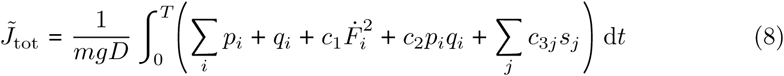

The final issue, poor scaling, was partially helped by normalization (next section), but further ameliorated through the automatic scaling feature (‘automatic-hybridUpdate’) in GPOPS-II [32]. Automatic scaling requires all unbounded variables to be redefined with finite bounds. Selected bounds are shown in S1 Table, and were chosen to be much larger than the expected magnitude of the variables, but small enough to improve scaling of the optimal control problem (see S2 Appendix for further details).

#### Normalization

All variables were normalized. Times were normalized to *T* ≡ *D/U*, lengths were normalized to *l*_*b*_, and forces were normalized to *mg*, yielding a nondimensional time constant

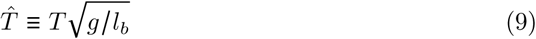

In this paper, normalized variables are noted with a prime exponent (*e.g. x* → *x*′. Since the force rate coefficient *c*_1_ (Eq 2) may have biological significance, it was also normalized. However, since 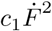 must have units of J s^−1^, the proper normalization is 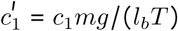, breaking dynamic similarity, and making *l_b_* and *m* an explicit part of the optimization. [25] noticed a similar issue with their rate penalty breaking dynamic similarity. Instead of the above normalization, we determined an appropriate normalized force-rate penalty constant 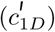 for the Dalmatian documented in [33] moving with time constant 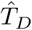. For all other cases, 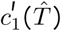 was scaled relative to 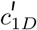 as

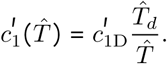

The augmentations to the cost function (Eq 8) were not scaled.

While the characteristic length for the computational problem was *l*_*b*_, we will report non-dimensional speed as the square root of the Froude number,

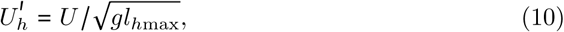

following a common convention in other literature [1, 33].

#### Implementation of the trajectory optimization problem

The optimal control problem was solved using a Direct Collocation (DC) method with *hp*-Adaptive Gaussian Quadrature, implemented in GPOPS-II (v. 2.1) [32]. This software also supplied mesh-refinement methods. The resulting nonlinear problem was solved using the nlp-solver SNOPT (v. 7.4-1.1) [34, 35]. GPOPS-II mesh tolerance was set to 10^−4^.

Numerical solutions to optimal control problems require an initial guess. Values at each of the 16 initial grid points for each variable were chosen from a uniform pseudo-random distribution within the bounds of each variable. An exception is the horizontal position state variable, which was set as [0, *D*] for the start and ending conditions, respectively, and interpolated linearly between those values for intermediate grid points.

The algorithm for finding the pseudo-global optimum (the optimum among all valid solutions) is as follows. An initial randomized guess was passed to GPOPS-II with one mesh refinement step. For this initial step, the slack variable orthogonality constraint (Eq 4) was explicitly enforced with *c*_2_ and all *c*_3*i*_ set to 0 (Eq 8). This step allowed the program to more quickly approach a feasible region. The output was then down-sampled and passed back to GPOPS-II, this time with two mesh iterations steps, no slack variable orthogonality constraint and with the fully augmented objective (Eq 8, with *c*_2_ = 1 E - 3 and *c*_3*i*_ = [100 10 10] (relaxation parameter coefficients for P3, P4 and P5, respectively). This process was repeated two more times, with *c*_2_ = 1 E - 3 and *c*_3*i*_ = [1000 100 100], with three and finally 8 mesh iterations. The final solution was saved and the problem reset with a new random guess, until the required number of random guesses were explored. Other settings used for SNOPT and GPOPS-II are listed in S2 Table.

For exploring the effect of *c*_1_ on the solution in one test case, 100 initial guesses were used. For the three detailed test cases, 500 initial guesses were used. For comparison to the large empirical dataset from [36], the number of guesses depended on the speed, since lower speeds required longer solution times per guess but the final solutions were not strongly affected by the initial guess. For 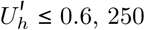, 250 guesses were used; for 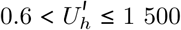 500 guesses were used; for 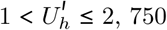, 750 guesses were used; and for 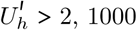, 1000 initial guesses were used.

Each solution was determined as “valid” on the basis of (1) satisfactory mesh error (lower than the tolerance of 1e-4) and (2) satisfactory (low) constraint violation. The second criteria was especially critical for complementarity conditions, which were not explicitly enforced and were violated more often than the other constraints.

Among valid solutions, the lowest cost solution was selected as the (pseudo-)global optimum as determined by *J*_tot_ (Eq 3). That is, since constraints were checked through the above validation step, the augmented parts of the objective (regarding constraint violation) were ignored as part of the relative optimality of one solution as compared to the others.

#### Empirical data

The model requires five inputs: stride length, speed, body length, center of mass position, and limb lengths. As validation of the model, duty factor, phase offset, and ground reaction force profiles are also required. These empirical data were acquired for single speeds from four studies [33, 37, 38], representing dogs moving at a slow walk, medium walk, and trot, respectively. In addition, [36] provides curves for duty factor, stride length, and phase offset as a function of speed from a large dataset. While all these studies supply a length calibration (*e.g.* shoulder height, withers height), they do not supply all necessary body proportions (except for [33], which supplies shoulder and hip locations in Figure 12 of that study). These additional proportions were taken from sagittal view images of the breed standard from the American Kennel Club website [39–41]. The acquired parameters are summarized in Table 1.

**Table 1.** Empirical data used for validation and as input (bold) for the model. PL is pair lag, the fraction of stride between left hind and left fore touchdown (sometimes referred to as pair offset or phase offset). DFh and DFf are mean duty factor of the hind and fore limbs, respectively. *Not reported at the original source [36]; derived from [37]

| Main Source: | [37] | [33] | [38] | [36] |
| --- | --- | --- | --- | --- |
| Breed | Mix: 50%<br>Rhodesian<br>Ridgeback | Dalmation | Labrador<br>Retriever | Belgian<br>Malinois |
| $m'_f$ ($m'$) | **0.63** (0.95) | **0.61** (1.12) | **0.66** (1.01) | **0.63\*** |
| $l'_h, l'_f$ | **0.91, 0.75** | **0.89, 0.79** | **0.86, 0.72** | **1.01, 0.92** |
| $l_b$ (m) | 0.63 | 0.61 | 0.52 | 0.48 |
| Proportions<br>source: | [41] | Fig 12 in [33] | [40] | [39] |
| $U'_h, D'$ | **0.34, 1.3** | **0.39, 1.21** | **1.15, 2.0** | $D' = 1.04 + 1.36U'_h$ |
| Hildebrand gait<br>classification | Walk in diagonal<br>couplets lateral<br>sequence | Walk in singlefoot<br>lateral sequence | Running trot | Variable |
| <u>Gait Formula</u> |  |  |  |  |
| PL | 0.15 | 0.20 | 0.50 | $U'_h$ Dependent |
| DFh | 0.65 | 0.61 | 0.40 | " |
| DFf | 0.69 | 0.67 | 0.51 | " |

Forelimb length was taken as the length from manus to shoulder joint in standing. Hindlimb length was taken as length from pes to hip joint in standing. Body length was taken as hip to shoulder joint in standing. Masses were assumed concentrated at shoulder and hip, following [37]. To determine center of mass location (or, equivalently, the anterio-posterior body mass distribution [42]), a balance of torque approach was used following [43]. Briefly, since the net moment about the center of mass must be zero during steady, periodic locomotion, the net torque produced by the hindlimbs in the sagittal plane must be equal and opposite to the forelimbs. Assuming the trunk is approximately horizontal during the stride, then

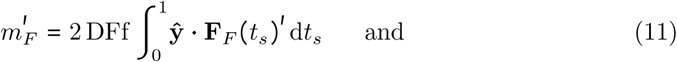

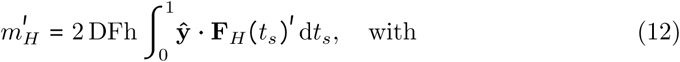

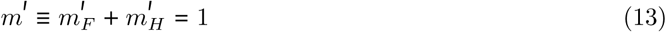

where DFf and DFh are the duty factors of the fore and hind limbs, respectively, **ŷ** is the vertical unit vector, **F**_*F*_ and **F**_*H*_ are the fore and hindlimb ground reaction forces, and *t*_*s*_ is “stance time” (time during stance for a given foot, normalized to stance duration). 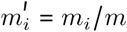 where *m* is the total body mass. Due to measurement error, the final condition is not always met. To correct for these errors, 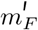 and 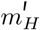 were scaled by 1/*m*^′^, ensuring that their sum was exactly 1 (Table 1).

Forces were derived from Fig 8 in [37], while the footfall sequence was derived from Fig 9. Ground reaction forces and footfall sequence were also pulled from Fig. 5a in [33], and Fig 1 in [38] and Fig 3 in [44], for comparison to modelled predictions.

**Fig 3.**
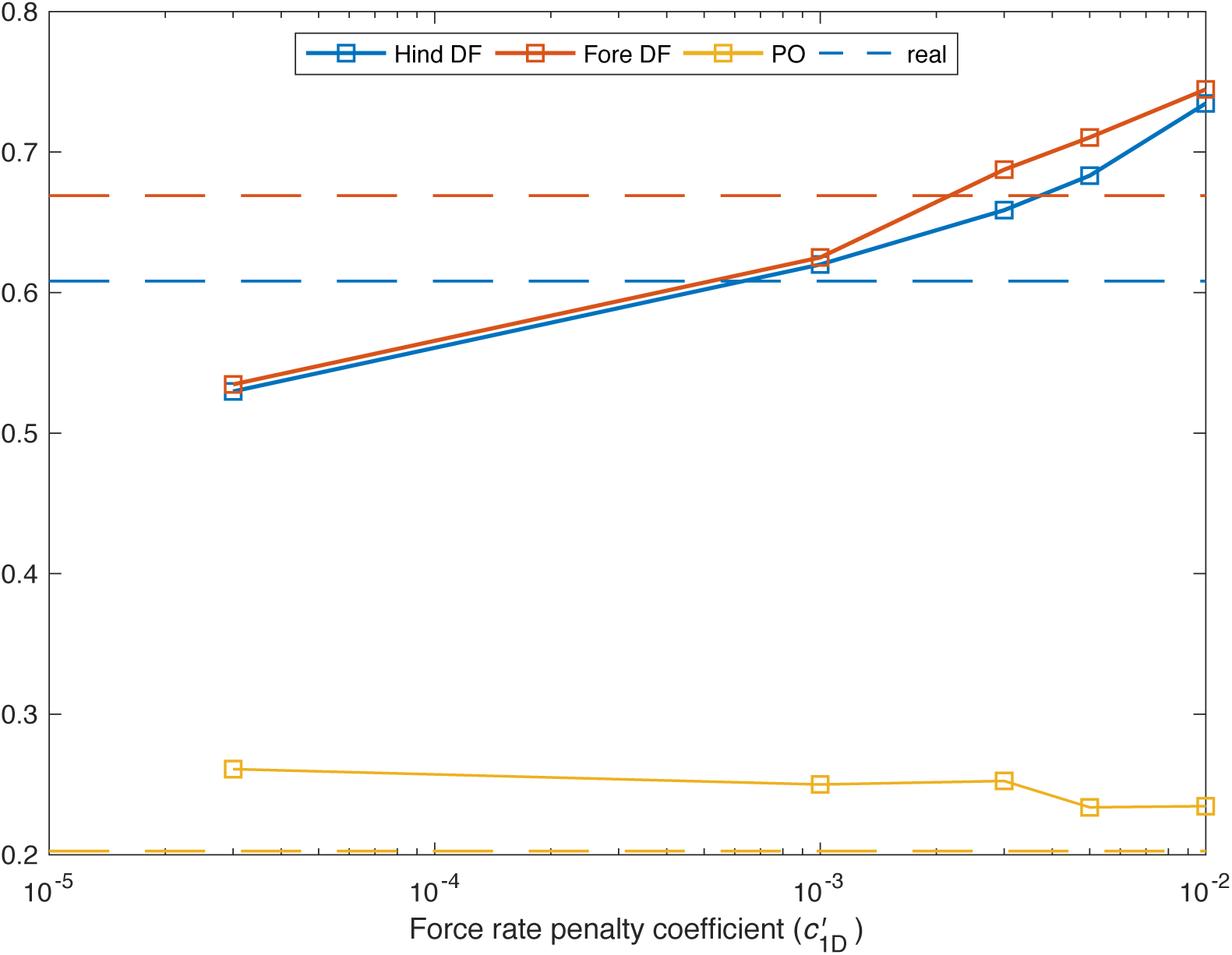
Optimal duty factors and left-fore phase offsets (pair lag) *vs* magnitude of force rate cost coefficient 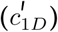 at 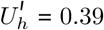. As force-rate penalties increase, duty factor increases to slow the generation and reduction of force. Optimal duty factors remain close, but exhibit some divergence at larger 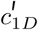. Phase offset decreases slowly with force rate penalty. At no point do optimal duty factors and phase offset simultaneously match the empirical values (dotted lines) from [33]. The morphological data is based on a Dalmation from [33] (see also Fig 4B).

The standard error in mean phase offset from [37] was found by the authors to be small, and so was not reported for most limbs. Variation in footfalls could not be calculated from [33] as these authors reported only one footfall sequence at the speed of interest. The standard deviation in duty factor was reported by [38] in their Table 1, but error in phase offset was not reported. For this, we took the range of phase offsets observed in Fig 1 in [38] as a measure of the natural variation in phase offset. We used the same approach for both phase offset and duty factor in [44], using their Fig 3b. Error bars in gait diagrams for our Fig 4 therefore represent twice standard deviation where available, and range of observed values otherwise.

**Fig 4.**
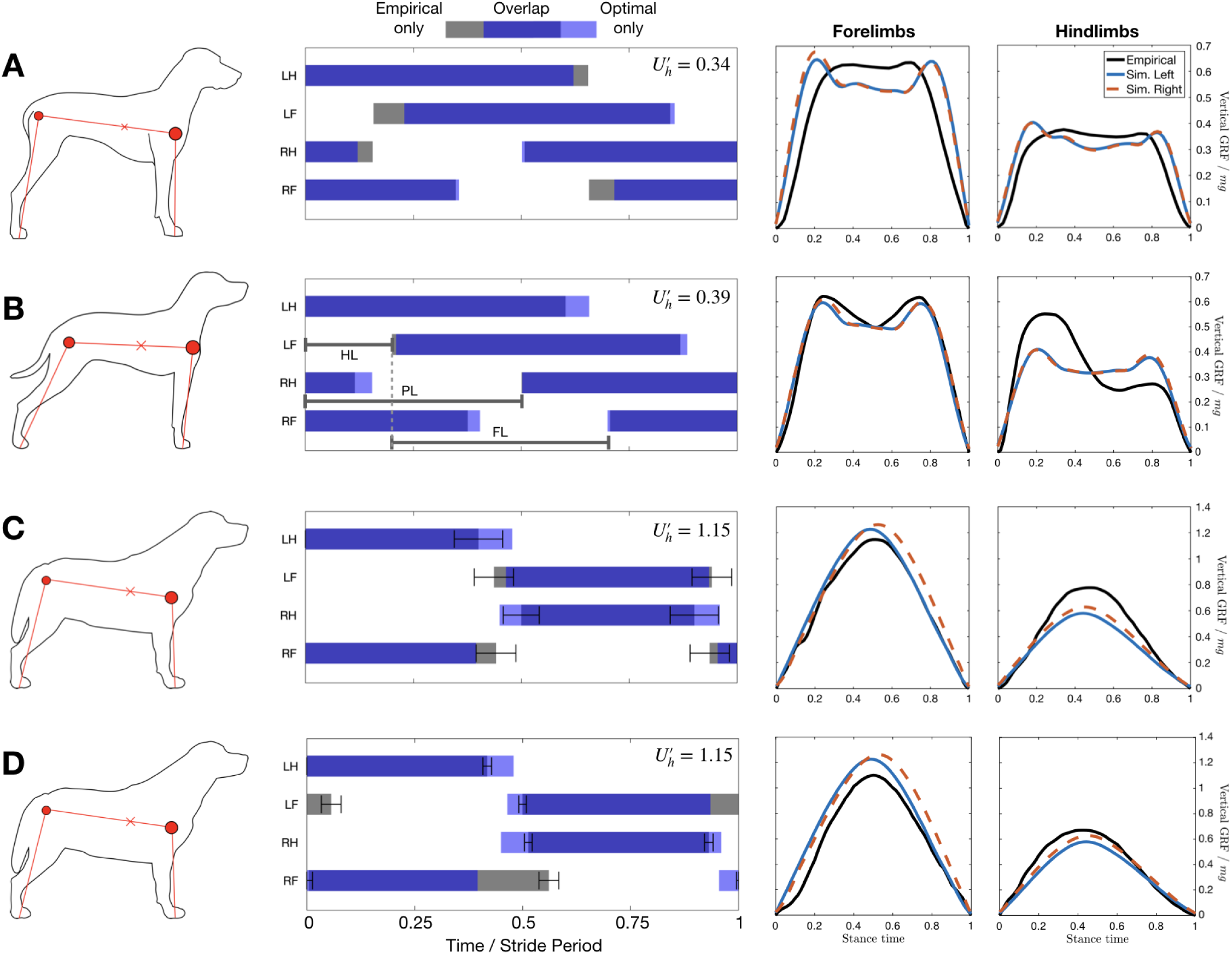
Optimal solutions compared to empirical data for three test cases. Left column shows an outline of the dog breed in question, and the points used as morphological measurements. Middle column shows that gait diagrams for the empirical (grey) and optimal solutions (light blue) have substantial overlap (dark blue). Right two columns demonstrate that empirical ground reaction forces (GRF, black lines) qualitatively match optimization results. The optimization GRF from left (solid blue line) and right limbs (dashed red line) are very similar, reflecting the symmetrical solutions that are discovered. At slow speeds (A-B), the simulation discovers a singlefoot walking gait, while at an intermediate speeds (C-D) it discovers trotting, matching natural gait choice in dogs. All optimizations use 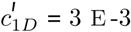. Empirical data and morphological measurements from [37, 41] (A), [33] (B), [38, 40] (C) and [40, 44] (D). For left column, circles show shoulder and hip positions, with relative size indicating the relative mass carried at these points. x indicates centre of mass position. For middle column, LH = Left Hind limb, RH = Right Hind, LF = Left Fore, RF = Right Fore. Error bars in phase are described in the methods.

We required empirical gait transition speeds to compared to our model. The concept of “a transition speed” in dogs is somewhat misleading, and Maes *et al.* [36], observed more of a transition zone: speeds where dogs were almost equally likely to be choosing one gait or another. We quantified the probability of dogs choosing one gait or another using their data (Fig 4 in [36]), and binned observations in 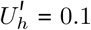 increments. Trotting and Pacing were combined as one gait, as were Transverse, Slow Rotary and Fast Rotary Gallops. The relative frequency (number of observations for a particular gait divided by the total number of observations at that speed) are shown in S1 Fig. We defined the gait transition point as the speed where the relative frequency of one gait (*e.g.* walking) transitions from greater than 0.5 to less than 0.5.

#### Data availability

Data supporting this work is available as supplemental files using a comma-delimited format.

## Results and Discussion

### Initial validation and fitting of rate cost

The simulation was run for the morphology, speed and stride length derived from [33], as outlined in Table 1 for a number of rate penalties (*c*_1_ in Eq 2). Empirically, the prescribed speed corresponded to a moderate walk [9]. The simulation spontaneously discovered a four-beat walk– specifically, a singlefoot walk [9]– as cost-minimizing regardless of rate penalty.

For all rate penalties, the simulation predicts a “double-hump” shape in ground reaction force to be optimal (Fig 2), a common characteristic of both bipedal and quadrupedal walking [45]. In bipedal locomotion, this shape arises from the pre-heelstrike pushoff [46–48], a dynamic “trick” for minimizing energetic losses at foot contact. The pre-heelstrike pushoff redirects the center-of-mass velocity so that it is less in line with the leading limb, thus reducing the amount of negative work the leading limb needs to do [24, 49]. The same “trick” seems to be discovered by the model; a second peak in the trailing limb ground reaction force occurs prior to the first peak in the leading limb.

The amplitudes of the peaks are affected by the magnitude of the force-rate cost. A purely work-based objective is expected to have impulsive foot-down and push-off [18, 19, 29, 30], but this would incur a large force rate penalty. The model compromises on a solution with smooth GRF, with larger peaks if the force rate penalty is small, and smaller peaks if the penalty is large.

Five multipliers on rate penalties are compared in Fig 3. As rate penalty increases, both fore and hind limb duty factors increase and remain relatively similar, but exhibit increasing divergence. Duty factor is strongly dependent on rate penalty because mean vertical force must remain constant (supporting weight), but longer actuation time allows the required force to be supplied over a longer time. At the lowest rate penalties, duty factor and phase offset approach the work minimizing solutions of 0.5 and ≈ 0.25, respectively, for symmetrical walking [50]. Phase offset is fairly insensitive to rate penalty, but exhibits a slight decrease with increased force-rate cost.

While either fore or hind duty factor can be tuned to be arbitrarily close to empirical values, matching both is not possible through adjusting rate penalty alone in this case. Furthermore, since phase offset is relatively insensitive to rate penalties, it is likewise difficult to tune. The results of this analysis imply that, while adding a rate penalty enhances agreement with empirical data, it alone cannot explain the discrepancy between work-minimizing solutions and empirical results, at least for this highly simplified model. This finding is in contrast to human locomotion models, where the simple addition of a force-rate penalty enhances empirical agreement substantially [24, 29].

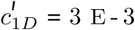 matches the empirical GRF profiles (Fig 2) and duty factors (Fig 3) for this test case fairly well. It is used as the coefficient of choice for further exploration in the remaining test cases.

### Detailed comparison to four test cases

Three cases are examined in detail in Fig 4. In the first two cases, the optimization correctly predicts a symmetrical, four-beat walking solution with double-hump ground reaction forces. In the third and fourth case, the optimization correctly predicts two-beat run with near-simultaneous fore-hind contact and single-hump GRFs. The optimization also correctly predicts the gait to be a “grounded run” [51], in the sense that it exhibits in phase potential and kinetic energy changes, but no flight phase (Fig 5H). For trotting, two empirical results from related studies [38, 44] are compared to the same optimization solution, demonstrating the range of “normal behaviour” in trotting for this breed. Overall, the simulation matches either empirical result fairly well, demonstrated by the substantial overlap of stance and similar shape of the GRFs, but matches neither perfectly.

**Fig 5.**
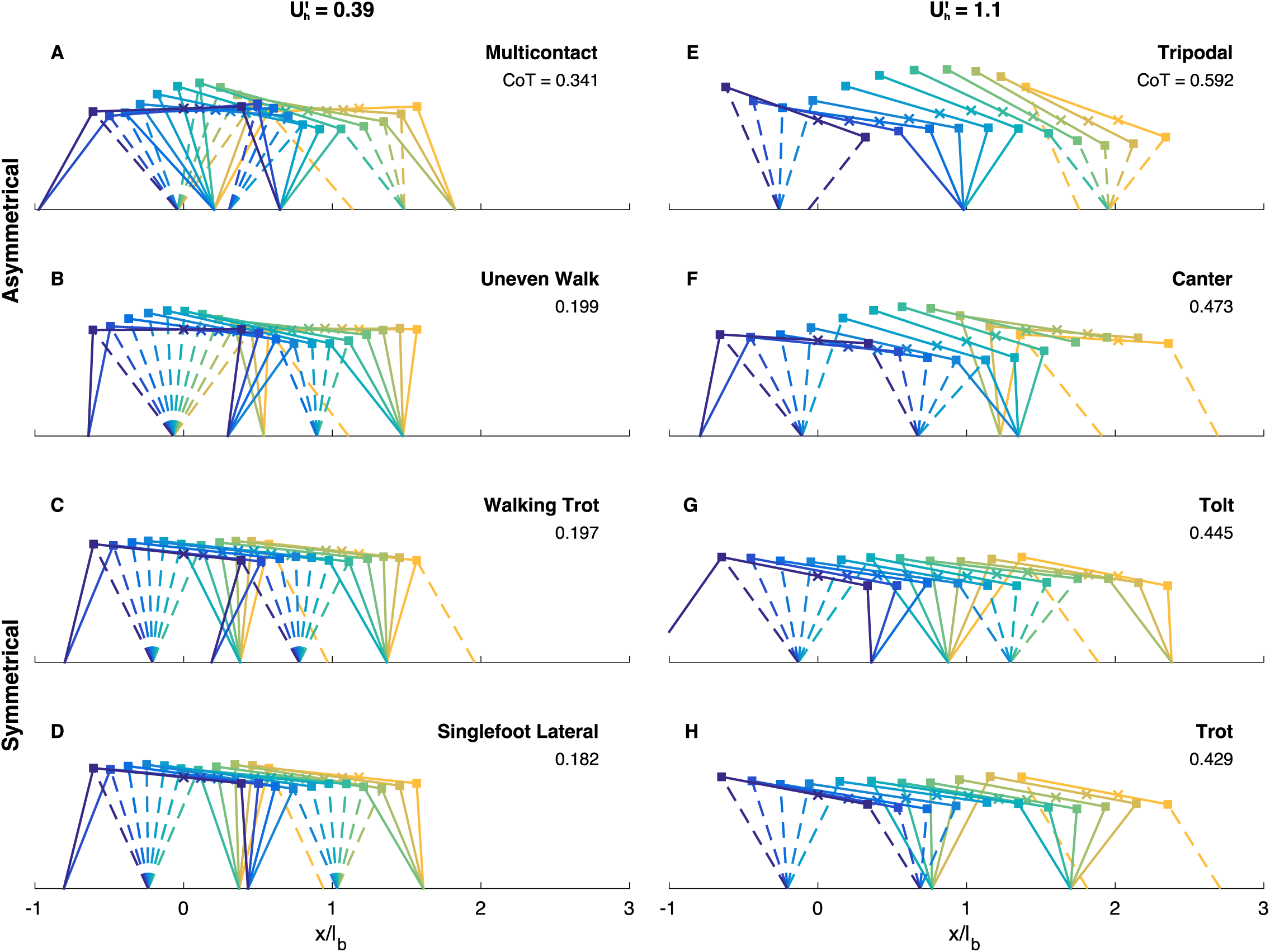
These locally optimal solutions are a sample of the diverse gaits possible with the model. Each stick figure is a freeze-frame of a different part of the gait cycle. The first still (leftmost, purple) is at touchdown of the left hind limb. Each still is separated from the next in time by one tenth of the stride time, culminating at the next left-hind touchdown. x marker represents center of mass location. At both a walking speed (left column) and running speed (right column) asymmetric (upper two rows) and symmetric gaits (bottom two rows) are possible. *y*-axis scale is equal to the *x*-axis.

The substantial overlap between optimization and empirical gait diagrams demonstrates the success of the optimization approach. It must be emphasized that any of the 5040 quadrupedal foot contact sequences could have been selected by the optimization. While, in a planar sense, many of these gaits are the same (for example, there’s no distinction between a lateral and diagonal sequence gait), there are still 2352 event sequences that are unique in a planar sense. Moreover, the model could even exclude limbs, adding more options to its repertoire.

Fig 5 shows some examples of locally optimal gaits discovered by the optimizer. These included asymmetrical (Fig 5A,B,E,F) and symmetrical gaits (Fig 5C,D,G,H), some that can be related to natural gaits (Fig 5C,D,F-H), and some that are clearly unnatural (Fig 5A,B). The model also did sometimes use fewer than four limbs (Fig 5E). Yet the pseudo-globally optimal gaits are readily identified as those naturally chosen by dogs at those speeds (Fig 5D,H) and match the empirical solutions fairly closely (Fig 4B-D).

Only the second case (Fig 4B) was “tuned” with the force rate penalty; the same penalty was used in all other cases. Yet even within this tuning, the optimization always selected a gait that was qualitatively similar to the empirical data (Figs 2 and 3). These results come despite fairly rough empirical estimates, varying morphological parameters, and many simplifying assumptions in the model (avoiding springs, most joints, and using the simplest possible mass distribution).

Forces are predicted fairly well, at least qualitatively in terms of shape and magnitude. While shapes of the ground reaction forces are an optimization decision, the relative magnitude of these forces are highly constrained. The mass distributions used were taken from empirical vertical forces in the first place (Eq 11–13), and kinematic periodicity ensures that net torque around the body is zero. Furthermore, mean net vertical force must equal body weight. Assuming the body remains relatively horizontal, then these two conditions completely determine the mean vertical force produced by either pair of limbs. While these arguments emphasize that little merit should be seen in matching relative magnitudes (apart from duty factor predictions), they do show that the quadruped’s center of mass must be biased towards one pair of limbs to get fore-hind differences in (mean) GRF magnitude. Without bias, there can be no difference in mean vertical forces produced by the fore and hind limbs; single-point mass models of quadrupedal locomotion cannot capture this effect [52].

While the mean magnitude of GRF is not very informative, the shape is. The optimization predicts a double-hump GRF profile for walking, and single hump for trotting. These predictions are born out in the empirical cases, although the magnitude of the force peaks in the slowest case is substantially lower than predicted. The empirical forces for the hind feet in the intermediate walking case (Fig 4B) were derived using a questionable method from simultaneous contact of fore and hind limbs on a force plate [33], and are not considered reliable.

The shape of simulation forces appear realistic for two reasons. First, the force-rate penalty ensures they are smooth (as averaged or filtered empirical ground reaction forces tend to be). Second, the work cost promotes double-hump walking and single hump trotting GRFs. The double-hump profile is representative of a “pre-heelstrike” trailing-limb pushoff, which redirects velocity prior to contact of the leading limb, avoiding some negative work. The roughly symmetrical, single hump profile is representative of a short, largely vertical contact of the limbs, which are work minimizing in point-mass bipedal models [49]. Our interpretation is in contrast to [16], who say all shape characteristics (double hump, single humps) are from “*oscillation modes*” due to springs and resonant properties of the system. Despite not having any springs, similar double-hump walking and single hump running GRFs emerge from our energy-minimizing model.

While spring-mass systems are often used to approximate the emergent gait behaviour, they are not the only system that describes the behaviour well and should not be taken as a prerequisite for “spring-like” behaviour. Rather, spring-like behaviour emerges from energy minimization [17–19, 49], even in the absence of springs.

It is difficult to tell if differences between solutions in Fig 4 are due to differences in morphological input or prescribed speed. To control for these differences, the model was compared to a large dataset using Belgian Malinois dogs of similar sizes over a large speed range [36].

### Comparison to a large kinematic dataset

The results of the simulations are compared to empirical results in Fig 6. Optimal mean duty factor (averaged across all limbs) matches empirical values very well for 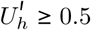, (Fig 6A) following its slow decay as speed increases. For 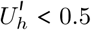, the predicted duty factor is almost constant at approximately 0.6. As with the other test cases (Fig 4), the model predicts singlefoot walking as the optimal gait at low speeds, and trotting as optimal at high speeds (Fig 6B). The phase offset is predicted well for all walking and trotting speeds, with all phase offsets lying within 0.1 of the empirical values for 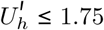. The model predicts a sharp transition to a trotting gait between 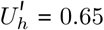 and 0.7, close to the empirical transition point of 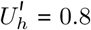 (S1 Fig). Curiously, the simulation chooses a transition speed almost exactly at the slowest self-selected trotting speed 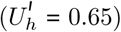 observed by Maes *et al.* [36] (S1 Fig). While a true gallop never emerges from the optimization, even well past the natural trot-gallop transition of 1.9 (S1 Fig), slight asymmetry begins to emerge from the solution for 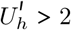 (Fig 6C).

**Fig 6.**
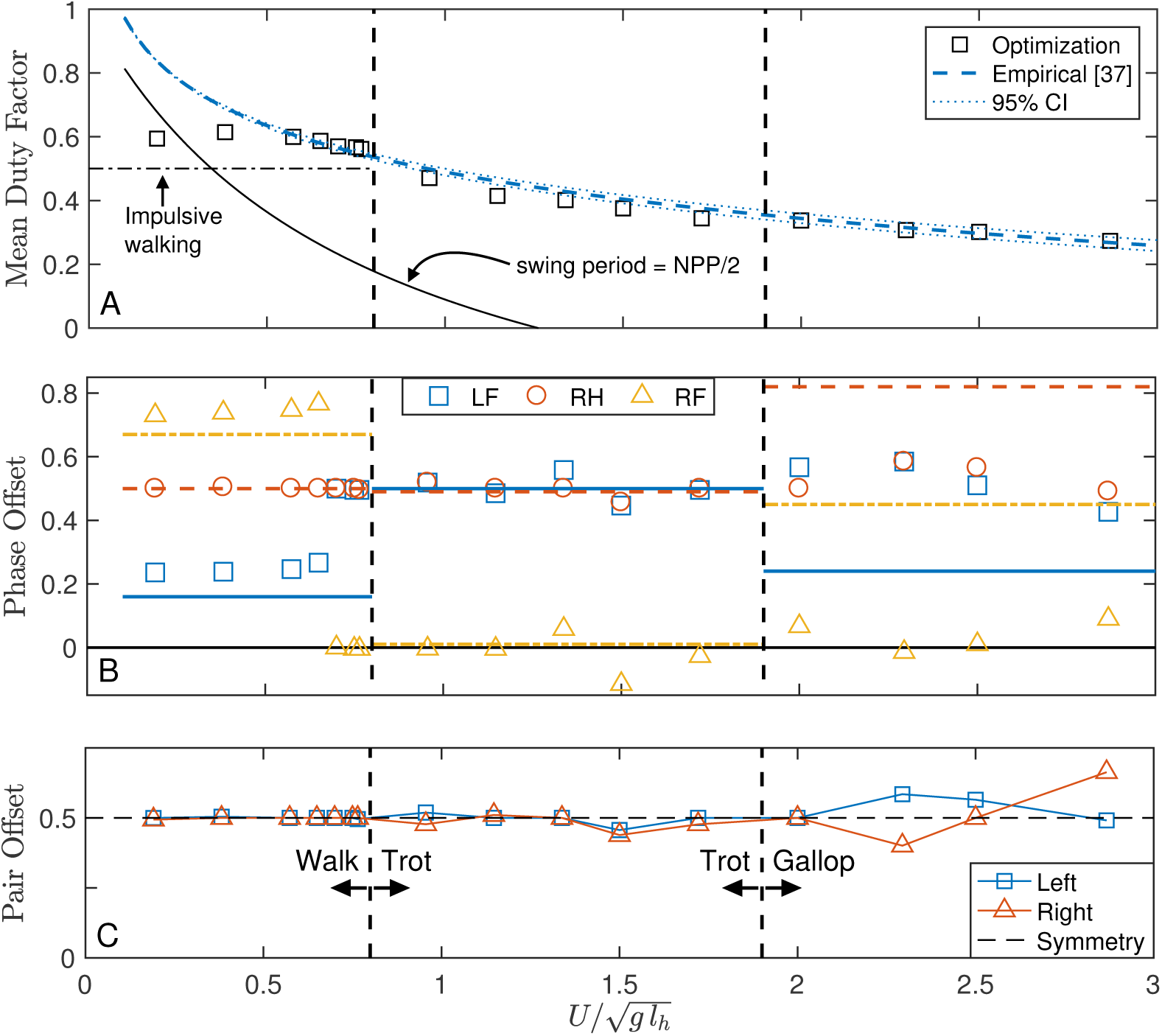
Pseudoglobal optimal solutions (markers) are compared to a large dataset for Belgian Malinois dogs [36]. (A) A slowly decaying duty factor with increasing speed is predicted well by the model for medium to fast speeds, using *c*_1*D*_ = 3 E - 3. At slow speeds, the model remains constant, slightly above impulsive predictions, while the empirical data approaches 1. A swing period at half the natural pendular period (NPP) does not explain the discrepancy. 95% CI for empirical duty factor is shown as a thin dotted line. (B) The optimal phase offset matches empirical values well for walking and trotting speeds, and the optimal walk-trot transition is very close to the natural transition speed. However, the model does not transition to a gallop after the natural trot-gallop transition speed (C) Pair offset (time between contact of the hind and fore limbs) is shown against speed. A perfectly symmetrical gait would have pair offsets of 0.5 in both left and right limbs; walking and trotting in dogs is highly symmetrical both naturally [53] and in the model. Galloping is an asymmetrical gait, and past the trot-gallop transition, the optimal gait becomes asymmetrical. Empirical phase offsets are mean values for each gait [36]

### Response of optimal duty factor to speed

Above 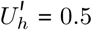, optimal values follow the empirical trendline closely. The decaying behaviour of duty factor is at odds with an impulsive trot (DF = 0), which would be work-optimal. Below about 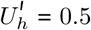, the optimal duty factor levels off at about 0.6, slightly above the work-minimizing prediction of 0.5 for a walk [50]. A similar pattern of constant duty factor shifting suddenly to a decay with increasing speed was also observed by Hubel and Usherwood [25] in their biped model with an analogous work + rate penalty objective (though the rate penalty was peak-power in their case, rather than force rate).

Work-minimal walking should have DF = 0.5 because double stance involves costly simultaneous positive and negative work [30, 48]. Work-minimal running gaits should have DF = 0 because a vertical impulse at stance avoids costly fore-aft decelerations. The increase in duty factor above work-minimizing predictions in our model is due to the force rate penalty, which allows both these additional costs in order to avoid impulses. However, as speed increases, enhancing stance duration involves ever-increasing excursion angles, and so larger and larger fore-aft forces, increasing work costs. Therefore, the optimal solution is to decrease duty factor as speed increases to manage this tradeoff.

It is interesting that the simple addition of a rate penalty yields such high agreement with duty factor at high speeds, especially since the model does not include massive limbs, springs, or a compliant back, nor does it use the correct (galloping) gait at high speeds. If rate penalties (whether force/time, power or activation penalties) are real phenomena, we would expect their influence to be most pronounced at higher speeds, where, since stride time is shorter, by necessity forces must be produced more quickly, power is higher, and muscles must be activated more frequently.

The discrepancy at lower speeds can be partially explained by leg swing dynamics. There is a cost to swinging legs during locomotion [54, 55] that is minimized when the leg is swung close to its natural frequency [56]. As a result, quadrupeds (including dogs) tend to decrease stance duration, rather than swing duration, as speed increases [36, 57].

If swing costs dominated, at all speeds it would be advantageous to the animal to adjust its duty factor such that swing time was exactly half the natural swing period, *i.e.*

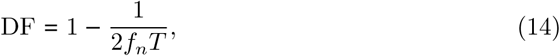

where *f*_*n*_ is the leg natural frequency and *T* is stride time. Taking *T* as given (based on empirical data), and using the allometric equations from [58] for dog leg natural frequencies, we can generate the predicted curve shown in Fig 6a. It takes on a similar shape to the empirical data at low speeds, but lies well under the curve. According to the data from [36], dogs drive their legs in swing at about twice natural frequency on average during walking. The reasons for this are unclear; nevertheless, including limb moment of inertia might account for some of the discrepancy between the predicted duty factor and empirical data.

### Response of Phase Offset to speed

Phase offset for each limb from the optimal solutions are compared to empirical data from [36] in Fig 6B. Left-hind touchdown (LHTD) is set at *t* = 0, and all phase offsets are timed relative to LHTD. For ease of comparison, at natural walking speeds optimal gaits are transformed so that, if left fore touchdown (LFTD) occurs after right fore touchdown (RFTD), the left and right forelimbs are swapped. For trotting and galloping speeds, the opposite transformation is used. Since the model is planar, these transformations result in no change in cost while making all two-beat gaits “trots” and all four-beat gaits lateral sequence. While in theory quadrupeds can perform 5040 event sequences, swapping left and right limbs leaves 2352 unique “planar” quadrupedal footfall sequences for the optimizer to choose from; any of these sequences can be performed at any speed, but only a small number are found to be optimal.

At low speeds, the model predicts a fore-hind phase difference between 0.23 and 0.27, close to the 0.25 phase predicted from impulsive walking with even mass distribution and equal fore and hindlimb lengths [50, 59]. The mean empirical phase offset for walking of 0.16 ± 0.05 (mean ± standard deviation) does not differ greatly from either of these values. For 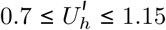, the optimal solution is to use a symmetric trot, with closely-associated fore-hind contacts. Above 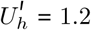, the fore-hind contact becomes dissociated, with forelimbs leading or lagging associated hindlimbs by 0.05 to 0.11. For 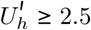, the gait becomes somewhat asymmetrical, with either the fore or hind pair exceeding a 0.5 phase difference. In no case did a galloping gait emerge as globally optimal, despite the model settling on galloping as locally optimal in a number of cases.

We find that galloping should not use less work than trotting for this model, in agreement with others who have analyzed alternative rigid back models [16, 60]. The difference in cost between galloping and trotting is slight, however. For example, the model spontaneously discovered a galloping solution with CoT = 0.58, only 1% higher cost than the minimal-cost trotting solution at the same high speed of 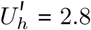 (CoT = 0.56). Previous studies have shown that galloping becomes optimal at high speeds with an elastic, compliant spine [61–64]. Future work could add active compliance to our model’s trunk, to see whether true elastic return is necessary for the optimality of galloping, or whether any form of compliance (even active compliance) makes galloping more economical.

Despite initializing the optimization from over 8000 randomized initial guesses, only four-beat walking and two-beat running (or slight deviations thereof) were discovered as pseudo-global optimal solutions. This is strong evidence (but not definitive proof) that these are globally optimal solutions for the quadrupedal configuration investigated (Fig 1) with a work-based cost and force-rate penalty.

Again, these results emerge despite the simplicity of the modelling approach, and the absence of springs. This is further evidence that natural walking, trotting and pacing are energetically optimal even without elastic elements. We are not suggesting that tendons (or springs in robotic quadrupeds) do not potentially reduce energetic cost, only that they are not necessary *prerequisites* for the optimality of natural gait. A gait could emerge as a recurrent strategy within a species because of its low energetic cost, and well-tuned tendons (such as the digital flexor and interosseus tendons in horses [65]) can then arise through natural selection to complement those strategies.

The discovery of trotting (as opposed to tolting) as energetically optimal is consistent with an analysis of Usherwood [66]. A dimensionless pitch moment of inertia can be defined as [67]

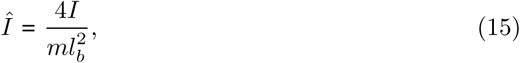

or the moment of inertia relative to the radius of gyration of the body. To a first approximation, for *Î* > 1, tolting is energetically optimal; otherwise trotting is [66]. In our case, *Î* = 0.93, not far from a more realistic estimate of 0.84 for dogs [33, 68]. Since these dimensionless moments of inertia are less than 1, we expect the model (and dogs) to find trotting to be less expensive than a running walk.

However, some deviation from exact trotting– perfect symmetry and fore-hind offset of 0.5– was observed at higher speeds. Moreover, while a clear singlefoot contact pattern was observed for 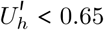, and a clear trotting pattern emerged for 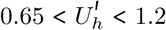, at higher speeds the footfall pattern was less consistent. Pair dissociation became more common, but was not always optimal (for example at 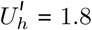 and 2.2) and the optimal gait became asymmetric (but regularly increasing asymmetry was not observed in the pseudoglobally optimal solutions; Fig 6C).

One reason for the lack of a consistent pattern may be that there are more variable locally optimal gaits at higher speeds. To explore this possibility, we plot phase offsets for all valid solutions for four speeds from the walking, trotting, slow gallop and fast gallop ranges (Fig 7). Each solution is both feasible (satisfying the constraints and dynamics), and locally optimal (that is, the NLP solver reported successful convergence). Each solution lives in a 3D space of foot contact phase lag: [Hind Lag, Pair Lag, Fore Lag] = [HL,PL,FL], defined as fraction of time between touchdown of left-hind and right-hind (Hind Lag), left-hind and left-front (Pair Lag) and left-front and right-front (Fore Lag). These phase lags are sketched in Fig 4B and follow the basic terminology used by Abourachid *et al.* [69]. Note that our Pair Lag (PL) differs from their definition (PL_Abourachid_ = 1 – PL), since we use the left-hind leg as reference while Abourachid *et al.* used right-front. Color represents cost of each solution, with dark blue as low cost, yellow as high cost, and light blue as intermediate.

**Fig 7.**
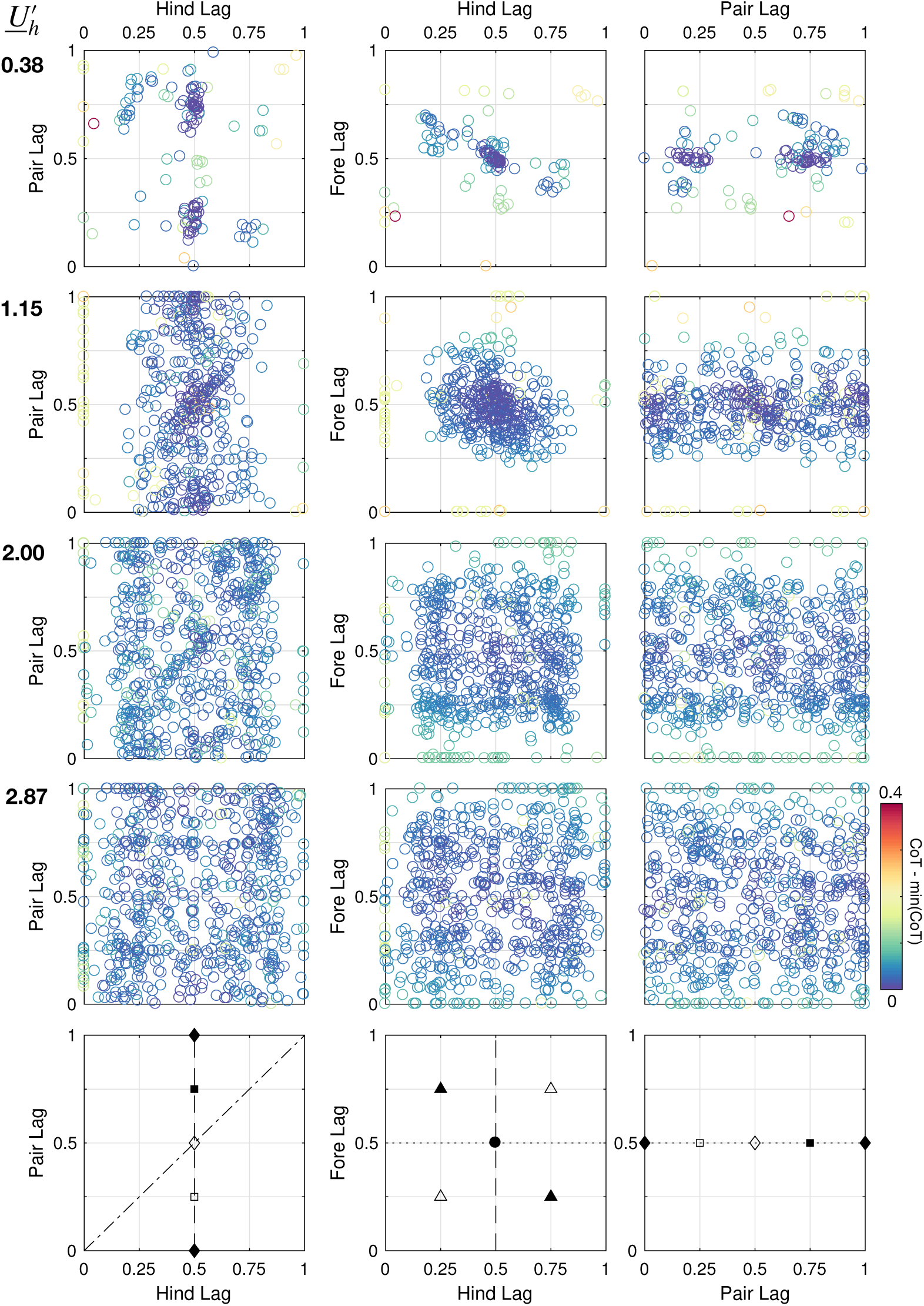
All local optima found for the model for each speed based on a Belgian Malinois morphology. Each row corresponds to a given speed condition, and each data point is a feasible local optimum arising from energy optimization from one random guess, with relative cost (CoT - min CoT for a given 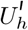) indicated by color. Solutions are plotted in terms of Hind Lag (HL), Pair Lag (PL) and Fore Lag (FL), defined in Fig 4. As speed increases, solutions become more variable. Pseudo-global optimal solutions for each case are shown in Fig 6. Bottom row: Some recognizable gaits and their positions in the plots. Dashed line: hind symmetry; dotted line: fore symmetry; dot-dash line: simultaneous right-hind and left-front contact (as in a canter) ▪ diagonal singlefoot ▫ lateral singlefoot ◊ trot ♦ pace ▵ transverse gallop ▴ rotary gallop • perfect symmetry

As speed increases, less solution clustering is observed. At a slow speed 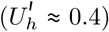, only two clusters emerge, at [0.5, 0.25, 0.5] and [0.5, 0.75, 0.5], representing symmetrical singlefoot walking (lateral or diagonal, respectively [9]). At a higher speed 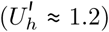, clustering becomes less pronounced, but two clusters emerge; one at [0.5, 0.5, 0.5] (symmetrical trotting) and another at [0.5, 0, 0.5] (symmetrical pacing). However, unlike the walking case, symmetry is not as clearly optimal, as represented by the large scatter of solutions between 0.25 < {HL, FL} < 0.75 compared to 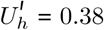

At a slow galloping speed 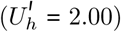, clustering has all but disappeared. Still, bands of viable solutions emerge; in particular, many solutions emerge in the range 0.20 < {HL, FL} < 0.80 with any PL, while solutions are rare outside this range. A wide band of low-cost solutions emerges at FL = 0.5, with some bias towards HL = 0.5 (symmetrical trotting). However, small clusters also emerge at the nodes [{0.25, 0.75},PL,{0.25, 0.75}], representing galloping (transverse if HL = FL, rotary otherwise). Interestingly, solutions cluster around the PL = HL line in the leftmost plot, representing simultaneous contact of fore and hind limbs. A canter would lie on this line.

Interestingly, small clusters of high-cost solutions also emerge at {HL, FL} = {0, 1}, representing a bound, half-bound, or pronk (depending on PL and the difference between HL and FL). These solutions emerged at all speeds, but were never globally optimal, reflecting the results of [16].

Why are solutions more variable at higher speeds? One reason may be that phase and event sequence impact cost less at higher speeds than at lower speeds. Fig 8A shows how cost changes with increasing speed. Reflecting both human [70] and animal studies [10], the cost of transport in walking is highly sensitive to speed, while the cost of transport of running is much less so; however, minimum and median cost always increases with speed. Despite the increased costs in running, the range in CoT is fairly small at higher speeds compared to walking. Relative to the increases in cost, the CoT range of feasible solutions decreases considerably from lowest speed to highest speed (Fig 8B) At high speeds, virtually all combination of phase are explored (Fig 7), with little change in cost (Fig 8). Optimizing locomotion at higher speeds appears to involve searching a flat and fuzzy cost landscape for the best gaits.

**Fig 8.**
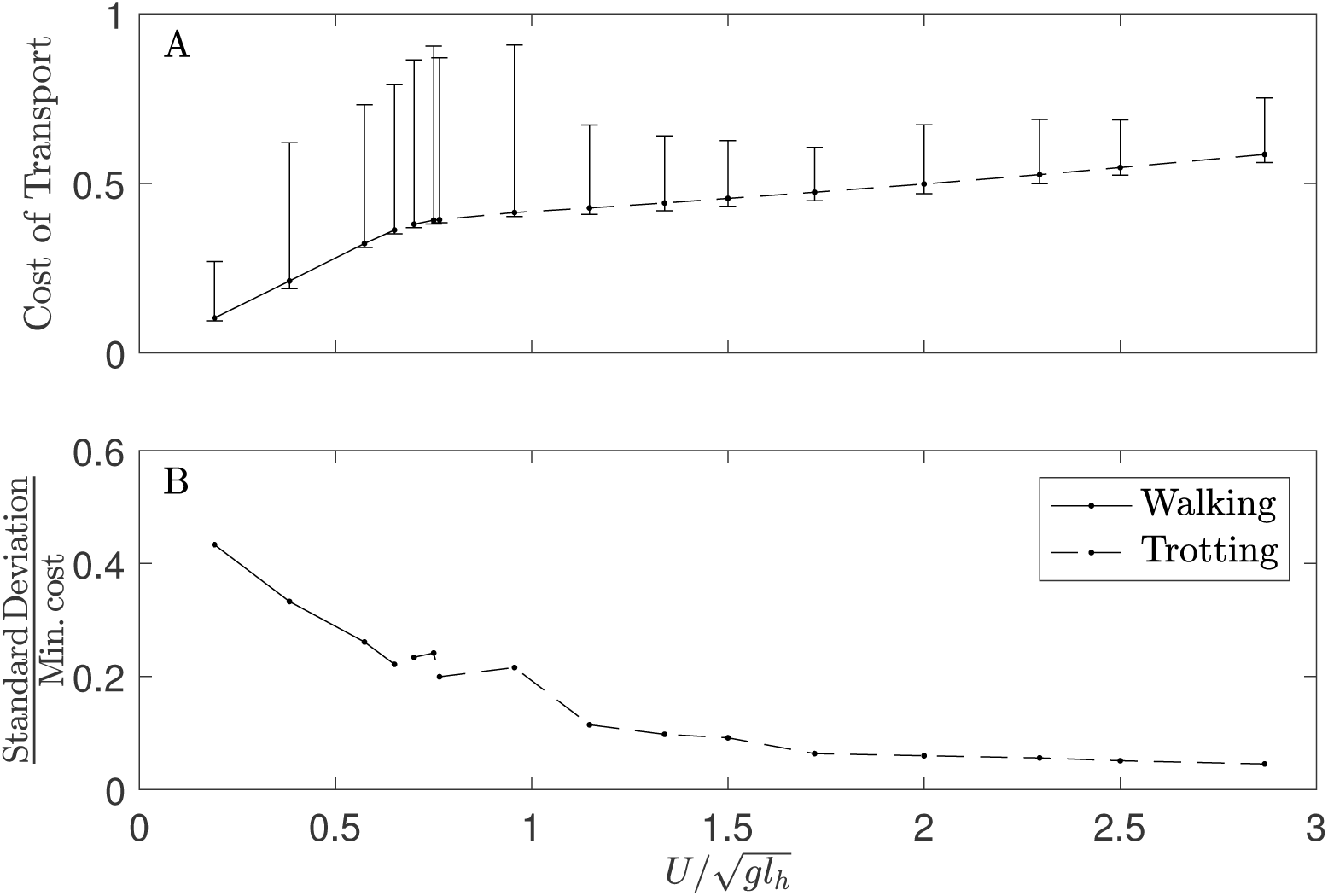
(A) Median Cost of Transport increases linearly for walking speeds (solid line), but exhibits a sharp change to a slower rate of increase after the walk-trot transition (dashed line), mirroring the response to speed of walking and running observed in human data [71]. The range of the costs of feasible solutions (whiskers) increases up to the walk-trot transition, and then settles into a near-constant range. The distribution of costs is heavily skewed, as indicated by the median value approaching the minimum at all speeds. This indicates that the solvers tended to discover solutions with costs close to the minimal value, but occasionally were “trapped” in local optima with costs far from the minimum. (B) As speed increases, the standard deviation of the distribution of costs of transport gets smaller, relative to the minimal cost, indicating that the variance in costs of local optima is relatively smaller at higher speeds than at lower speeds

The variability in locally optimal solutions may somewhat reflect reality; dogs exhibit substantial natural variation in gait at higher speeds. For example, empirical data from [38, 44] indicate that duty factor and pair dissociation can vary at the same trotting speed in labradors (Fig 4C,D). The transition from trot-gallop is much less pronounced than walk-trot (S1 Fig), and while trots and rotary gallops are the most common running gaits, dogs will also pace and use transverse gallops [36]. If the relative cost between these gaits are not substantial, then factors apart from energetics might dictate locomotory decisions.

The plots in Fig 8 represent a combination of symmetrical and asymmetrical solutions. Hildebrand [53] studied the symmetrical gaits of domestic dogs in great detail, and discovered that dogs (across a large range of speeds and breeds) choose a limited subset of symmetrical gaits from the total “gait space” available to them (every combination of duty factor and pair lag from 0 to 1). Does the limited scope of *C. lupus familiaris* gait represent a fluke of evolution, a physical constraint, or is it the result of an energy optimization process?

In Fig 9 we plot all the symmetrical gaits preformed by dogs and recorded by Hildebrand [53], as well as the “C-shaped” contour that he thought captured the scope of locomotory behaviour most meaningfully. Overlaid on these data are all the feasible convergent results from every condition studied in Fig 6. Color represents relative cost, that is the deviation of the cost for a given solution from the minimal cost at the same speed. Each data point is a complete optimization from a single random guess, resulting in a solution that satisfied all dynamics and constraints, and where successful convergence towards local optimality was indicated by the NLP solver.

**Fig 9.**
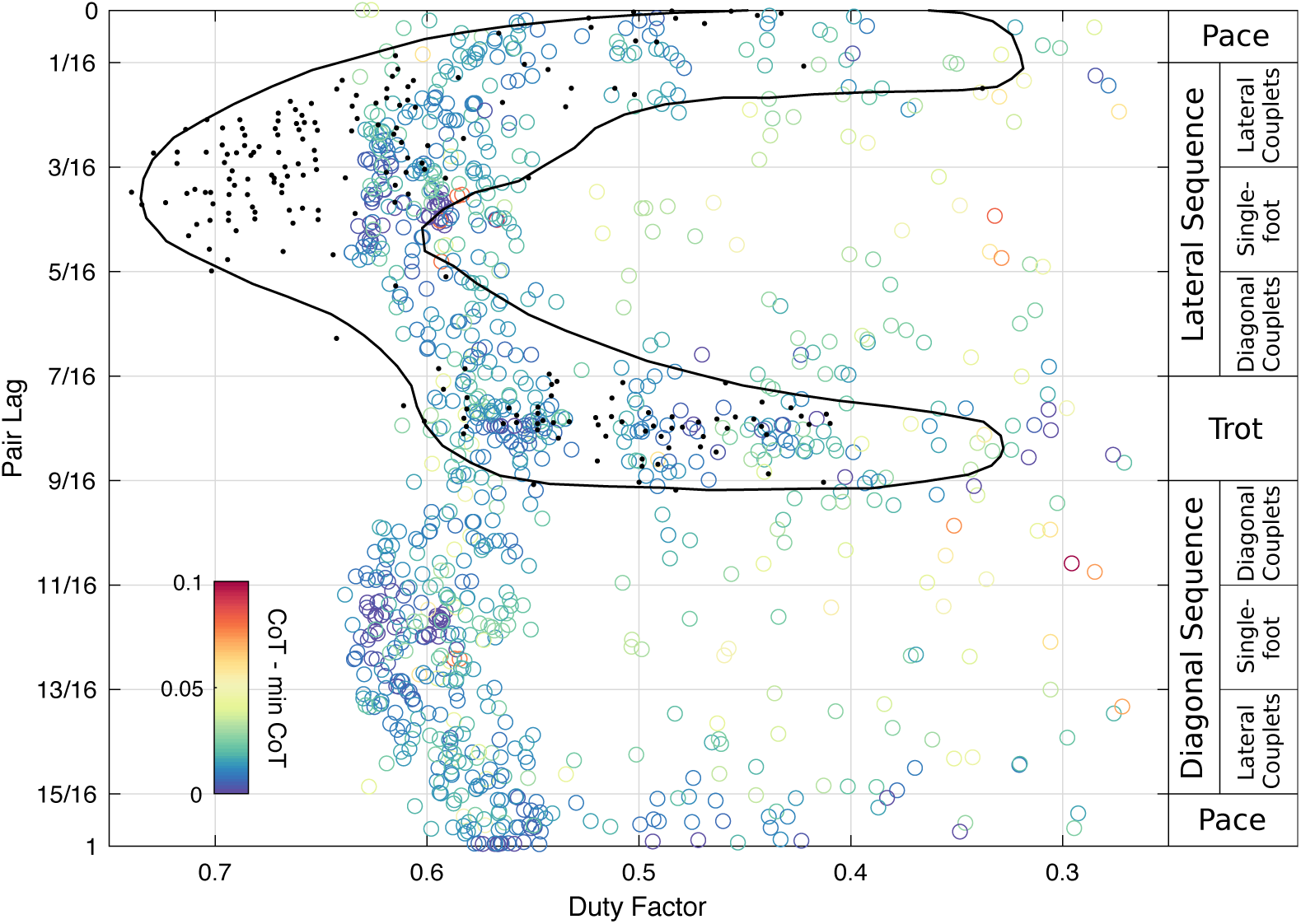
Symmetrical gaits discovered by the model compared to all symmetrical gaits in dogs observed by Hildebrand [53]. Data points (black) from Fig 1 and contour (line) in Fig 2 of [53], representing mean duty factor and Pair Lag for symmetrical gaits in dogs, are overlaid on all locally optimal symmetrical solutions (coloured circles) discovered by the model for the Belgian Malinois dataset (Fig 6). The empirical contour takes on a “C” shape in the upper (lateral sequence) region of the plot. The optimal solutions take on two “C” shapes, one in the upper half and one in the lower, as lateral and diagonal sequence gaits have equal cost in a planar model. While there is some discrepancy in lower duty factor, there is substantial overlap between Hildebrand’s contour and the clustering of locally optimal solutions in the model. Each coloured data point represents one solution from a different random guess at any of the speeds represented in Fig 6. Colour represents cost relative to the minimal cost solution at that given speed. Gait definitions, according to Hildebrand [9], are shown on right.

The vast majority of symmetrical solutions, as well as the lowest cost, lie within a similar “C-shape” to that observed by Hildebrand. Infrequently, sub-optimal but feasible solutions are discovered outside the range observed by Hildebrand. Their presence indicates that such solutions are possible– so there seems to be no plausible physical constraint to prevent dogs from entering that region of gait space– but their relative high cost indicates that they are unlikely to be chosen from an optimization process.

In contrast, the solutions along the C contour are frequently discovered by the optimization process. All are local optima; solutions that the optimizer converged on. The remarkable similarity between the distribution of optimal solutions and Hildebrand’s empirical observations suggests that dogs are also optimizing cost of transport when selecting gait.

Notably, Hildebrand’s results combined observations from dog breeds of varying sizes and shapes (from Basset hounds to Great Danes [53]), yet the simulations used only one morphological set, based on Belgian Malinois. The consistency between simulations and empirical data, despite the differences in size, is due to dynamic similarity. While a small and large dog may move differently at the same absolute speed, they are moving at different dimensionless speeds (the square root of Froude numbers, Eq 10) and so have different dynamic constraints and opportunities. Yet when their dimensionless speed is similar, their behaviour is similar [1]. As our simulation probed a large range of Froude numbers, we explored the scope of dynamic regimes that dogs (of all *sizes*) might be expected to experience in level, steady movement.

Harder to explain is the similarity between our results and Hildebrand’s despite differences in morphological *shape*. The similarity seems to suggest that general shape (at least as far as limb length to body length ratios and mass distribution) has little effect on the optimality of trotting, pacing, and single foot walking over other symmetrical gaits. The effect of morphology on the optimality of gait is something that could be explored with the present model by systematically varying morphological inputs.

One major discrepancy between the empirical and simulation results is the presence in the latter of a lower “C” that is a reflection of the first, upper “C”. In this planar model there is no energetic distinction between a pair lag of *p* and *p* ± 0.5, so it should come as no surprise that for every optimal gait in the upper region, there is an equally optimal gait in the lower region with the same duty factor but with a phase increase of 0.5. However, the discrepancy opens the question of why dogs (and most other quadrupedal mammals [7, 72]) use symmetrical gaits with pair lag less than 0.6 as opposed to higher pair lags. This is a question that a planar model cannot answer, but may be explained by considering full three-dimensional dynamics.

## Conclusions

We developed a minimally constrained quadrupedal model capable of any of the 2300+ planar contact sequences available to quadrupeds. Of those possibilities, it found two basic gaits– four-beat walking in singlefoot, and two-beat trotting or pacing– as energetically optimal. It also transitioned spontaneously from walking to trotting at a realistic speed. While limb work was the chief contributor to cost, an additional force-rate penalty led to better prediction of duty factor. Despite no enforcement of gait symmetry, the optimal gaits were highly symmetrical except at high speeds, as observed in natural gait.

The GRF profiles were double-hump in walking and single-hump in trotting despite no springs being present in the model. This sheds doubt on the claim that these features result from “oscillation modes of the legged system’s compliance” [16]. Instead, they are the result of work-minimizing strategies; the “pre-heelstrike” pushoff in walking and near-vertical “pseudo-elastic” contact in running [49].

Galloping did not emerge as optimal at high speeds in our model, but elastic-limb models exhibit the same result if elastic elements are not included in the trunk. Future work will examine whether galloping emerges as optimal if the torso is allowed to extend or contract actively. This would test whether elastic elements of the spine are truly necessary for the optimality of galloping.

Limb work with a small force-rate penalty has substantial explanatory power. Despite simplified morphological parameters, the model predicted gait well at slow and intermediate speeds across several breeds of dog, demonstrating that detailed morphological modelling are not necessary to explain gait choice. While the factors affecting duty factor choice at slow walking remain a mystery, we believe that duty factor at higher speed is determined by the competing costs of work (lowering duty factor) and force rate (increasing it). However, little is known about the physiological mechanisms for this rate cost, and it could be an exciting avenue for further research.

Singlefoot walking and trotting emerged as optimal gaits despite over 8000 random guesses being used to seed optimization. This strongly suggests (but does not prove) that these gaits are globally optimal strategies at their respective speeds, at least for the modelling configuration used. It is unlikely that alternate gaits exist which would further minimize quadrupedal cost of transport at slow and intermediate speeds.

During locomotion, mammals are likely optimizing a work-based cost function with some form of force-rate related energetic penalty. Although elastic elements can lower metabolic cost of transport, they were likely not prerequisites to the evolution of modern gait in mammals, but rather complemented preexisting, metabolically optimal strategies.

## Acknowledgments

The authors would like to thank S. Javad Hasaneini for assistance in designing the trajectory optimization model, and Jessica M. Theodor (University of Calgary) for the use of her computing resources.

## 1 Supporting Information

**S1 Appendix. Mathematical statements of constraints.**

**S2 Appendix. Justification for variable bounds.**

**S1 Table. Optimization variable bounds.**

**S2 Table. Main settings for the optimization procedure.**

**S1 Fig. Relative frequency of walking, trotting/pacing and galloping as speed increases.** Walking shown in blue, trotting/pacing in red and galloping in green. Observations normalized to total count in each bin. Data from Maes et al., Fig 4 [36].

**S1 File. Optimal solutions for specific cases.** Pseudo-globally optimal ground reaction forces, footfall positions and initial kinematic conditions are given for the results with varying force-rate penalty, as well as the other test cases, as shown in Figs 2–4.

**S2 File. Duty factor, phase and cost for all valid solutions discovered for a range of speeds using Belgian Malinois morphology, as displayed in Figs 6–9**. ‘nlpinfo’ are the SNOPT exit conditions, with nlpinfo < 10 indicating satisfactory convergence on a local optimum. Phase is relative to the left-hind limb touchdown.

## References

1. Alexander RM, Jayes AS. A Dynamic Similarity Hypothesis for the Gaits of Quadrupedal Mammals. J Zool. 1983;201(1):135–152. doi:10.1111/j.1469-7998.1983.tb04266.x.

2. Alexander RM. Optimization and Gaits in the Locomotion of Vertebrates. Physiol Rev. 1989;69(4):1199–1227.

3. Biancardi CM, Minetti AE. Biomechanical Determinants of Transverse and Rotary Gallop in Cursorial Mammals. J Exp Biol. 2012;215(23):4144–4156.

4. Pennycuick CJ. On the Running of the Gnu (*Connochaetes Taurinus*) and Other Animals. J Exp Biol. 1975;63(3):775–799.

5. McGhee RB, Jain AK. Some Properties of Regularly Realizable Gait Matrices. Math Biosci. 1972;13(1):179–193. doi:http://dx.doi.org/10.1016/0025-5564(72)90033-8.

6. Alexander R, Langman V, Jayes A. Fast Locomotion of Some African Ungulates. J Zool. 1977;183(3):291–300.

7. Loscher DM. Kinematische Anpassungen Zur Kollisionsreduktion Im Schritt Vierfüßiger Lauftiere. Freie Universität Berlin; 2015.

8. Hildebrand M. Symmetrical Gaits of Horses. Science. 1965;150(3697):701–708.

9. Hildebrand M. Analysis of Tetrapod Gaits: General Considerations and Symmetrical Gaits. In: Herman RM, Grillner S, Stein P, Stuart D, editors. Neural Control of Locomotion. vol. 18 of Advances in Behavioral Biology. Plenum Press New York; 1976. p. 203–236.

10. Hoyt DF, Taylor CR. Gait and the Energetics of Locomotion in Horses. Nature. 1981;292:239–240.

11. Bertram JEA. Constrained Optimization in Human Walking: Cost Minimization and Gait Plasticity. J Exp Biol. 2005;208(6):979–991. doi:10.1242/jeb.01498.

12. Selinger JC, O’Connor SM, Wong JD, Donelan JM. Humans Can Continuously Optimize Energetic Cost during Walking. Curr Biol. 2015;25(18):2452–2456. doi:10.1016/j.cub.2015.08.016.

13. Polet DT, Schroeder RT, Bertram JEA. Reducing Gravity Takes the Bounce out of Running. J Exp Biol. 2018;221(3):jeb162024. doi:10.1242/jeb.162024.

14. Geyer H, Seyfarth A, Blickhan R. Compliant Leg Behaviour Explains Basic Dynamics of Walking and Running. Proc R Soc B Biol Sci. 2006;273(1603):2861–2867. doi:10.1098/rspb.2006.3637.

15. Gan Z, Wiestner T, Weishaupt MA, Waldern NM, Remy CD. Passive Dynamics Explain Quadrupedal Walking, Trotting, and Tölting. J Comput Nonlinear Dyn. 2016;11(2):021008.

16. Xi W, Yesilevskiy Y, Remy CD. Selecting Gaits for Economical Locomotion of Legged Robots. Int J Robot Res. 2016;35(9):1140–1154. doi:10.1177/0278364915612572.

17. Bertram JEA, Hasaneini SJ. Neglected Losses and Key Costs: Tracking the Energetics of Walking and Running. J Exp Biol. 2013;216(6):933–938. doi:10.1242/jeb.078543.

18. Srinivasan M, Ruina A. Computer Optimization of a Minimal Biped Model Discovers Walking and Running. Nature. 2006;439(7072):72–75. doi:10.1038/nature04113.

19. Hasaneini SJ, Macnab C, Bertram JEA, Leung H. The Dynamic Optimization Approach to Locomotion Dynamics: Human-like Gaits from a Minimally-Constrained Biped Model. Adv Robot. 2013;27(11):845–859.

20. Russ DW, Elliott MA, Vandenborne K, Walter GA, Binder-Macleod SA. Metabolic Costs of Isometric Force Generation and Maintenance of Human Skeletal Muscle. Am J Physiol-Endocrinol Metab. 2002;282(2):E448–E457. doi:10.1152/ajpendo.00285.2001.

21. Doke J, Kuo AD. Energetic Cost of Producing Cyclic Muscle Force, Rather than Work, to Swing the Human Leg. J Exp Biol. 2007;210(13):2390–2398. doi:10.1242/jeb.02782.

22. Dean JC, Kuo AD. Energetic Costs of Producing Muscle Work and Force in a Cyclical Human Bouncing Task. Journal of Applied Physiology. 2011;110(4):873–880. doi:10.1152/japplphysiol.00505.2010.

23. Kram R, Taylor CR. Energetics of Running: A New Perspective. Nature. 1990;346:265–267.

24. Kuo AD. A Simple Model of Bipedal Walking Predicts the Preferred Speed–Step Length Relationship. J Biomech Eng. 2001;123(3):264. doi:10.1115/1.1372322.

25. Hubel TY, Usherwood JR. Children and Adults Minimise Activated Muscle Volume by Selecting Gait Parameters That Balance Gross Mechanical Power and Work Demands. J Exp Biol. 2015;218(18):2830–2839. doi:10.1242/jeb.122135.

26. Kelly M. An Introduction to Trajectory Optimization: How to Do Your Own Direct Collocation. SIAM Rev. 2017;59(4):849–904. doi:10.1137/16M1062569.

27. Betts JT. Practical Methods for Optimal Control and Estimation Using Nonlinear Programming. Smith RC, editor. Advances in Design and Control. Society for Industrial and Applied Mathematics; 2010.

28. Croft JL, Schroeder RT, Bertram JEA. Determinants of Optimal Leg Use Strategy: Horizontal to Vertical Transition in the Parkour Wall Climb. J Exp Biol. 2019;222(1):jeb190983. doi:10.1242/jeb.190983.

29. Rebula JR, Kuo AD. The Cost of Leg Forces in Bipedal Locomotion: A Simple Optimization Study. PLOS ONE. 2015;10(2):e0117384. doi:10.1371/journal.pone.0117384.

30. Srinivasan M. Fifteen Observations on the Structure of Energy-Minimizing Gaits in Many Simple Biped Models. J R Soc Interface. 2010;8(54):74–98. doi:10.1098/rsif.2009.0544.

31. Baumrucker BT, Renfro JG, Biegler LT. MPEC Problem Formulations and Solution Strategies with Chemical Engineering Applications. Computers & Chemical Engineering. 2008;32(12):2903–2913. doi:10.1016/j.compchemeng.2008.02.010.

32. Patterson MA, Rao AV. GPOPS-II: A MATLAB Software for Solving Multiple-Phase Optimal Control Problems Using Hp-Adaptive Gaussian Quadrature Collocation Methods and Sparse Nonlinear Programming. ACM Trans Math Softw. 2013;41(1):1.

33. Jayes A, Alexander R. Mechanics of Locomotion of Dogs (*Canis Familiaris*) and Sheep (*Ovis Aries*). J Zool. 1978;185(3):289–308.

34. Gill PE, Murray W, Saunders MA. SNOPT: An SQP Algorithm for Large-Scale Constrained Optimization. SIAM Rev. 2005;47(1):99–131.

35. Gill PE, Murray W, Saunders MA, Wong E. User’ s Guide for SNOPT 7.7: Software for Large-Scale Nonlinear Programming. La Jolla, CA: Department of Mathematics, University of California, San Diego; 2018. CCoM 18–1.

36. Maes LD, Herbin M, Hackert R, Bels VL, Abourachid A. Steady Locomotion in Dogs: Temporal and Associated Spatial Coordination Patterns and the Effect of Speed. J Exp Biol. 2008;211(1):138–149. doi:10.1242/jeb.008243.

37. Griffin TM, Main RP, Farley CT. Biomechanics of Quadrupedal Walking: How Do Four-Legged Animals Achieve Inverted Pendulum-like Movements? J Exp Biol. 2004;2072(20):3545–3558. doi:10.1242/jeb.01177.

38. Bertram JEA, Lee DV, Case HN, Todhunter RJ. Comparison of the Trotting Gaits of Labrador Retrievers and Greyhounds. Am J Vet Res. 2000;61(7):832–838. doi:10.2460/ajvr.2000.61.832.

39. American Kennel Club. Belgian Malinois Dog Breed Information - American Kennel Club; 2017. http://www.akc.org/dog-breeds/belgian-malinois/.

40. Club AK. Labrador Retriever Dog Breed Information; 2018. https://www.akc.org/dog-breeds/labrador-retriever/.

41. Club AK. Rhodesian Ridgeback Dog Breed Information; 2018. https://www.akc.org/dog-breeds/rhodesian-ridgeback/.

42. Lee DV, Stakebake EF, Walter RM, Carrier DR. Effects of Mass Distribution on the Mechanics of Level Trotting in Dogs. J Exp Biol. 2004;207(10):1715–1728. doi:10.1242/jeb.00947.

43. Gray J. Studies in the Mechanics of the Tetrapod Skeleton. J Exp Biol. 1944;20(2):88–116.

44. Bertram JEA, Lee DV, Todhunter RJ, Foels WS, Williams AJ, Lust G. Multiple Force Platform Analysis of the Canine Trot: A New Approach to Assessing Basic Characteristics of Locomotion. Vet Comp Orthop Traumatol. 1997;10(3):160–169. doi:10.1055/s-0038-1632588.

45. Bertram JEA. Considering Gaits: Descriptive Approaches. In: Understanding Mammalian Locomotion: Concepts and Applications. John Wiley & Sons; 2016. p. 27–50.

46. Tucker VA. The Energetic Cost of Moving About. AmSci. 1975;63(4):413–419.

47. McGeer T. Dynamics and Control of Bipedal Locomotion. Journal of Theoretical Biology. 1993;163(3):277–314. doi:10.1006/jtbi.1993.1121.

48. Donelan JM, Kram R, Kuo AD. Simultaneous Positive and Negative External Mechanical Work in Human Walking. J Biomech. 2002;35(1):117 – 124. doi:http://dx.doi.org/10.1016/S0021-9290(01)00169-5.

49. Ruina A, Bertram JEA, Srinivasan M. A Collisional Model of the Energetic Cost of Support Work Qualitatively Explains Leg Sequencing in Walking and Galloping, Pseudo-Elastic Leg Behavior in Running and the Walk-to-Run Transition. J Theor Biol. 2005;237(2):170–192. doi:10.1016/j.jtbi.2005.04.004.

50. Usherwood JR, Williams SB, Wilson AM. Mechanics of Dog Walking Compared with a Passive, Stiff-Limbed, 4-Bar Linkage Model, and Their Collisional Implications. J Exp Biol. 2007;210(3):533–540. doi:10.1242/jeb.02647.

51. Usherwood JR, Szymanek KL, Daley MA. Compass Gait Mechanics Account for Top Walking Speeds in Ducks and Humans. J Exp Biol. 2008;211(23):3744–3749. doi:10.1242/jeb.023416.

52. Usherwood JR, Self Davies ZT. Work Minimization Accounts for Footfall Phasing in Slow Quadrupedal Gaits. eLife. 2017;6(5):630–650. doi:10.7554/eLife.29495.

53. Hildebrand M. Symmetrical Gaits of Dogs in Relation to Body Build. JMorph. 1968;124(3):353–359.

54. Marsh RL, Ellerby DJ, Carr JA, Henry HT, Buchanan CI. Partitioning the Energetics of Walking and Running: Swinging the Limbs Is Expensive. Science. 2004;303(5654):80–83. doi:10.1126/science.1090704.

55. Steudel K. The Work and Energetic Cost of Locomotion. I. The Effects of Limb Mass Distribution in Quadrupeds. J Exp Biol. 1990;154(1):273–285.

56. Doke J, Donelan JM, Kuo AD. Mechanics and Energetics of Swinging the Human Leg. J Exp Biol. 2005;208(3):439–445. doi:10.1242/jeb.01408.

57. Kilbourne BM, Hoffman LC. Scale Effects between Body Size and Limb Design in Quadrupedal Mammals. PLoS ONE. 2013;8(11):e78392. doi:10.1371/journal.pone.0078392.

58. Myers MJ, Steudel K. Morphological Conservation of Limb Natural Pendular Period in the Domestic Dog (Canis Familiaris): Implications for Locomotor Energetics. J Morphol. 1997;234(2):183–196. doi:10.1002/(SICI)1097-4687(199711)234:2%A1;183::AID-JMOR5%BF;3.0.CO;2-D.

59. Smith AC, Berkemeier MD. Passive Dynamic Quadrupedal Walking. In: Robotics and Automation, 1997. Proceedings., 1997 IEEE International Conference On. vol. 1; 1997. p. 34–39 vol.1.

60. Alexander RM, Jayes AS, Ker RF. Estimates of Energy Cost for Quadrupedal Running Gaits. J Zool. 1980;190(2):155–192.

61. Alexander RM, Dimery NJ, Ker RF. Elastic Structures in the Back and Their Role in Galloping in Some Mammals. J Zool. 1985;207(4):467–482. doi:10.1111/j.1469-7998.1985.tb04944.x.

62. Alexander RM. Why Mammals Gallop. AmZool. 1988;28(1):237–245.

63. Cao Q, Poulakakis I. On the Energetics of Quadrupedal Running: Predicting the Metabolic Cost of Transport via a Flexible-Torso Model. BB. 2015;10(5):056008.

64. Yesilevskiy Y, Yang W, Remy CD. Spine Morphology and Energetics: How Principles from Nature Apply to Robotics. Bioinspir Biomim. 2018;13(3):036002. doi:10.1088/1748-3190/aaaa9e.

65. Wentink GH. Biokinetical Analysis of the Movements of the Pelvic Limb of the Horse and the Role of the Muscles in the Walk and the Trot. AE. 1978;152(3):261–272. doi:10.1007/BF00350524.

66. Usherwood JR. An Extension of the Collisional Model of the Energetic Cost of Support Qualitatively Explains Trotting and the Trot-Canter Transition. J Exp Zool Part A. 2019; p. Forthcoming.

67. Raibert MH. Legged Robots That Balance. MIT Press; 1986.

68. Alexander RM. Optimum Walking Techniques for Quadrupeds and Bipeds. J Zool. 1980;192(1):97–117. doi:10.1111/j.1469-7998.1980.tb04222.x.

69. Abourachid A, Herbin M, Hackert R, Maes L, Martin V. Experimental Study of Coordination Patterns during Unsteady Locomotion in Mammals. J Exp Biol. 2007;210(2):366–372. doi:10.1242/jeb.02632.

70. Farley CT, McMahon TA. Energetics of Walking and Running: Insights from Simulated Reduced-Gravity Experiments. J Appl Physiol. 1992;73(6):2709–2712.

71. Hasaneini SJ, Schroeder RT, Bertram JEA, Ruina A. The Converse Effects of Speed and Gravity on the Energetics of Walking and Running. bioRxiv. 2017; p. 201319. doi:10.1101/201319.

72. Lemelin P, Schmitt D, Cartmill M. Footfall Patterns and Interlimb Co-Ordination in Opossums (Family Didelphidae): Evidence for the Evolution of Diagonal-Sequence Walking Gaits in Primates. J Zool. 2003;260(4):423–429. doi:10.1017/S0952836903003856.

