## Supplemental Appendix 1 for "A simple model of a quadruped discovers single-foot walking and trotting as energy optimal strategies"

### S1 Appendix: Mathematical Statements of Constraint

Five parameters are derived from empirical measurements, and given in the model. These are the stride length $D$, the stride period $T$, the body mass $m$, forequarter relative mass $m'_F = m_F/m$, body length $l_b$ We normalize lengths by the shoulder-to-hip length $l_b$, time by the stride period $T$, masses by the total mass $m$ and forces by the weight $mg$. The result is a normalized time constant

$$\hat{T} \equiv T\sqrt{\frac{l_b}{g}}$$

which appears in a number of equations. The vectors from the centre of mass to the fore and hindquarters are given, respectively, as

$$\mathbf{r}'_F = m'_H \left(\cos\theta \sin\theta\right), \quad \mathbf{r}'_H = -m'_F \left(\cos\theta \sin\theta\right).$$

Since the trunk is a rigid link with two point masses, the normalized moment of inertia is simply $I' = m'_F \cdot m_H$. Forces point along the legs, from the foot contact position $f'_{ij}$ to its associated articulation point (the shoulders or hips). Leading limbs act by definition through the forward contact positions $f'_{ij} + D'$. The limb length vectors are therefore

$$\begin{aligned} \mathbf{l}'_{ijT} &= \mathbf{x}' + \mathbf{r}'_i - f'_{ij}\hat{\mathbf{x}} \\ \mathbf{l}'_{ijL} &= \mathbf{x}' + \mathbf{r}'_i - (f'_{ij} + D)\hat{\mathbf{x}}, \end{aligned}$$

And the limb forces are

$$\mathbf{F'}_{ijk}(t) = F'_{ijk}\frac{\mathbf{l'}_{ijk}}{l'_{ijk}}.$$

As the body consists of only one rigid link, the dynamics are simply

$$\ddot{\mathbf{x}}' = (-\hat{\mathbf{y}} + \sum \mathbf{F}'_{ijk})\hat{T}^2 \quad (1)$$

$$\ddot{\boldsymbol{\theta}} = \frac{\sum \mathbf{r}'_i \times \mathbf{F}'_{ijk}}{I'}\hat{T}^2 \quad (2)$$

**Table A1.** A list of symbols used in this section and their meanings.

| Symbol | Meaning | Symbol | Meaning |
|---|---|---|---|
| $\hat{\mathbf{x}}$, $\hat{\mathbf{y}}$, $\hat{\mathbf{z}}$ | Fore-aft, vertical and right unit vectors | $F'_{ijk}$ | Limb force |
| $i = \{F, H\}$ | Fore, Hind | | Empirically Derived |
| $j = \{R, L\}$ | Right, Left | $l_b$ | Body length (m) |
| $k = \{T, L\}$ | Trailing, Leading | $l'_{F\max}$ | Shoulder to manus length in standing (body lengths) |
| $(\cdot)'$ | Normalized variable | $l'_{H\max}$ | Hip to pes length in standing (body lengths) |
| $\mathbf{x}$ | CoM position | $m$ | Total body mass (kg) |
| $\theta$ | Trunk pitch angle | $m'_F$ | Forequarter mass proportion |
| $f_{ij}$ | Footfall postions | $T$ | Stride period (s) |
| $p_{ijk}$, $q_{ijk}$ | Slack Variables ) | $D$ | Stride length (m) |
| $s_{\mathrm{a}ijk}$, $s_{\mathrm{b}ij}$, $s_{\mathrm{c}ijk}$ | Relaxation Parameters | | |

The normalized cost of transport is

$$J = \frac{1}{D'} \int_0^1 \sum_{ijk} \left( c'_{1\mathrm{D}} (\dot{F}'_{ijk})^2 \hat{T}_D / \hat{T} + p'_{ijk} + q'_{ijk} + c_{2a} s_{\mathrm{a}ijk} + c_{2c} s_{\mathrm{c}ijk} \right) + \sum_{ij} c_{2b} s_{\mathrm{b}ij} \, \mathrm{d}t \quad (3)$$

The other Path, Endpoint, and Integral constraints are given below

**Path Constraints**

P2: Hips and shoulders above ground: $(\mathbf{x} + \mathbf{r}_i) \cdot \hat{\mathbf{y}} > 0$

P3: Limb lengths do not exceed prescribed values: $(l_{i\max} - l_{ijk}) F_{ijk} - s_{\mathrm{a}ijk} \geq 0$

P4: Leading limbs are active strictly after trailing limbs $F_{ijT} \int_0^t F_{ijL} \, \mathrm{d}t - s_{\mathrm{b}ij} = 0$

P5: Limbs are below the body: $\mathbf{r}'_F \times \mathbf{F}'_{Fjk} - s_{\mathrm{c}Fjk} \geq 0$

$-\mathbf{r}'_H \times \mathbf{F}'_{Hjk} - s_{\mathrm{c}Hjk} \geq 0$

P6: Functional of abs. given by slack vars: $\mathbf{F}'_{ijk} \cdot (\mathbf{x}' + \mathbf{r}') - p'_{ijk} + q'_{ijk} = 0$

P7: Positive and negative parts of power are orthogonal:* $p'_{ijk} \cdot q'_{ijk} = 0$

*Only enforced in the first major iteration

**Endpoint Constraints**

E1: Kinematic periodicity: $$[y', \theta, \dot{x}', \dot{y}', \dot{\theta}]^{t'=0} - [y', \theta, \dot{x}', \dot{y}', \dot{\theta}]^{t'=1} = 0$$

E2: Continuity of forces: $$\sum_k F'_{ijk}(1) - F'_{ijk}(0) = 0$$

**Integral Constraints:**

I1: Equal Left-Right impulse: $$\int_0^T \sum_{i,k} \mathbf{F}'_{iLk} - \sum_{i,k} \mathbf{F}'_{iRk} \, \mathrm{d}t = 0$$
