## Supplemental Table 1 for "A simple model of a quadruped discovers single-foot walking and trotting as energy optimal strategies"

### Variable Bounds

| | Initial | | Intermediate | | Final | |
|---|---|---|---|---|---|---|
| | **Lower** | **Upper** | **Lower** | **Upper** | **Lower** | **Upper** |
| $t'$ | 0 | 0 | 0 | 1 | 1 | 1 |
| **STATES** | | | | | | |
| $x'$ | 0 | 0 | $-D'$ | $2D'$ | $D'$ | $D'$ |
| $y'$ | 0 | $4\max(l_{i\max})$ | 0 | $4\max(l_{i\max})$ | 0 | $4\max(l_{i\max})$ |
| $\theta$ | $-\pi/2$ | $\pi/2$ | $-\pi/2$ | $\pi/2$ | $-\pi/2$ | $\pi/2$ |
| $\dot{x}', \dot{y}'$ | $-4D'$ | $4D'$ | $-4D'$ | $4D'$ | $-4D'$ | $4D'$ |
| $\dot{\theta}$ | -4 | 4 | -4 | 4 | -4 | 4 |
| $F'_{ijT}$ | 0 | 10 | 0 | 10 | 0 | 0 |
| $F'_{ijL}$ | 0 | 0 | 0 | 10 | 0 | 10 |
| $\int_0^{t'} F'_{ijL}\,\mathrm{d}t'$ | | 0 | 0 | 10 | 0 | 10 |
| **PARAMETERS** | | | | | | |
| $f'_{ij}$ | $-\max(l_{i\max}) - D'$ | $\max(l_{i\max}) + D'$ | - | - | - | - |
| **CONTROLS** | | | | | | |
| $\dot{F}_{ijk}$ | -100 | 100 | | | | |
| $p_{ijk}, q_{ijk}$ | 0 | 20 | 0 | 20 | 0 | 20 |
| $s_i$ | 0 | 20 | 0 | 20 | 0 | 20 |
