## Supplemental Appendix 2 for "A simple model of a quadruped discovers single-foot walking and trotting as energy optimal strategies"

### S2 Appendix: Justification for variable bounds

We must have $\sum_{i,j,k} \int_0^1 \hat{\mathbf{y}} \cdot \mathbf{F}'_{ijk}(t) dt' = 1$, so if $F' \leq 10$ and only one limb is in contact with the ground, its shortest possible duty factor is 1/10. If all four limbs are operational, the shortest possible duty factor is 1/40. Dogs are never observed to have DF < 0.2, so this force limitation is well above the empirical limit.

If $|\dot{x}|$, $|\dot{y}| \leq 4D'$, then the COM speed is allowed to be four times the average horizontal translation speed. If $|\dot{\theta}| \leq 4$, then the body could conceivably more than fully rotate in one gait cycle. The maximum possible power (represented by $p_{ijk}$ and $q_{ijk}$) is for the force to be maximal when speed at either the hips or shoulders is maximal. The speed at either joint could be as high as $\max(\dot{\theta}) + \sqrt{\max(\dot{x})^2 + \max(\dot{y})^2} = 4(1 + \sqrt{2}D')$. $D'$ was as high as 5, so in principle the instantaneous power could have been as high as 323 (much larger than the limit of 20 set).

However, increasing the bounds on $p$ and $q$ led to larger solve times, and in practice, the peak power was much lower. Peak power was on occasion saturated, but only for solutions with unacceptably large complementarity violations (in particular, with hips and/or shoulders that operated with excursion angles > 180°). Among valid solutions, the largest peak power was only about 7. While these smaller limits are unlikely to have overly constrained the solution space, future analysis could open the bounds on these variables, at the expense of larger solve times.

Similarly, the limit for $|\dot{F}_{ijk}|$ was 100, which was much larger than any observed valid solution. Among valid solutions, the largest $|\dot{F}_{ijk}|$ was about 52. Other complementarity slack variables ($s_{ijk}$) never exceeded 1 E-2 for valid solutions.

Footfall positions were bounded as $|f_{ij}| < \max(l_{i\max}) + D'$. Geometrically, footfalls at their set bounds would not allow actuation without exceeding limb length constraints, so these limits cannot be exceeded.
