## Supplemental Table 2 for "A simple model of a quadruped discovers single-foot walking and trotting as energy optimal strategies"

Main Algorithm Settings

| | | Major Iteration | | | | |
|---|---|---|---|---|---|---|
| | | **1** | **2** | **3** | **4** | |
| | **Input Guess** | Uniform Random Distribution | Iteration 1 output | Iteration 2 output | Interation 3 output | |
| | ***pq* Complementarity** | Enforced by Constraint | Augmented objective (1e-3) | Augmented objective (1e-3) | Augmented objective (1e-3) | |
| | **Relaxation Parameter Coefficients** | [0 0 0]<br>[Limb Length, Trail-Lead, Limbs below] | [100 10 10] | [1000 100 100] | [1000 100 100] | |
| **SNOPT** | **Max Iter** | 500 | 2000 | 2000 | 2000 | |
| | **Tol** | 1e-06 | 1e-08 | 1e-08 | 1e-08 | |
| **MESH** | **Max Iter** | 1 | 2 | 3 | 8 | |
| | **Other settings (all major iterations)** | AutoScaling: Automatic-hybrid update | Method: RPM-integration | Derivatives: Sparse CD | Mesh Tol: 1e-4 | Initial mesh: 4 mesh intervals with 4 collocation points each |
