## Supplementary figures and images for "A simple model of a quadruped discovers single-foot walking and trotting as energy optimal strategies"

### Supplemental Figure 1

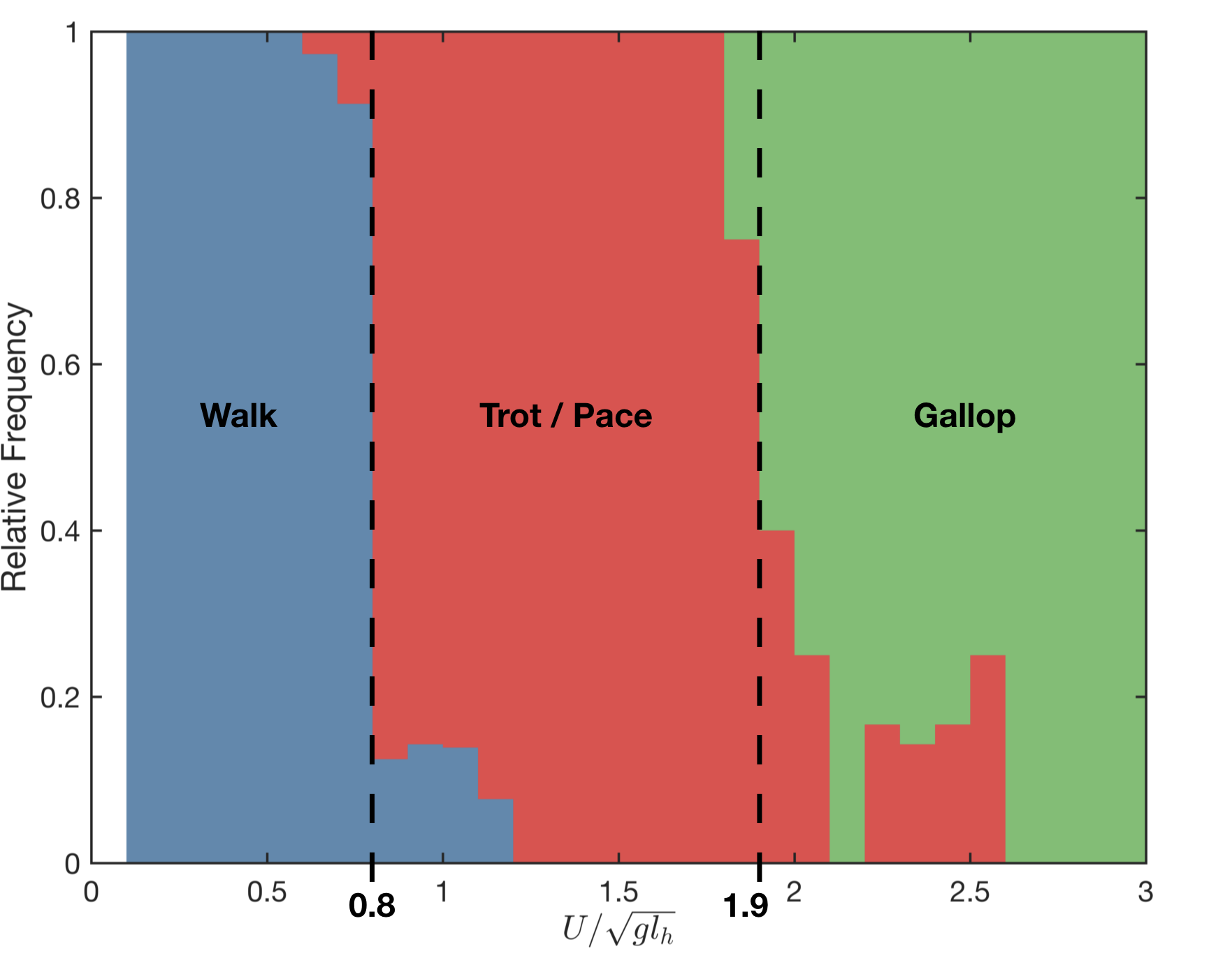
